# Folate-dependent one-carbon metabolism controls meiotic and post-meiotic epigenome remodeling in the male germline

**DOI:** 10.64898/2026.04.23.720283

**Authors:** Ai Ikuyo, Nozomu Fuse, Masaru Mori, Akiyoshi Hirayama, Yuto Yamada, Tatsuya Nakamura, Tomoko Sagi, Kai Otsuka, Satoshi H. Namekawa, Tomoyoshi Soga, Yohei Hayashi, So Maezawa

## Abstract

Environmental exposures can influence offspring health through epigenetic alterations in the male germline. Folate deficiency, a dietary perturbation that disrupts one-carbon metabolism and S-adenosylmethionine (SAM) production, has been linked to altered histone methylation and developmental abnormalities in offspring. However, when and how folate availability shapes the germline epigenome during spermatogenesis remains unclear. In this study, unbiased metabolomic profiling of spermatogenic cells uncovers stage-specific metabolic remodeling, including downregulation of serine– glycine–one-carbon (SGOC) metabolism in meiotic spermatocytes. Using a post-weaning folate-deficient mouse model, we investigate how folate availability influences germline epigenome establishment during spermatogenesis. Consistent with this metabolic transition, genome-wide chromatin accessibility profiling demonstrates extensive, stage-dependent remodeling under folate-deficient conditions, particularly in meiotic spermatocytes and post-meiotic spermatids. These accessibility changes display cell-type–specific genomic distributions and preferential localization to repressive chromatin compartments in post-meiotic cells. Histone modification analyses further reveal bidirectional redistribution of the active histone mark H3K4me3 in round spermatids. Although genome-wide distribution of the repressive mark H3K27me3 remains largely stable, folate deficiency alters its nuclear organization. Notably, a subset of H3K4me3 alterations established in post-meiotic cells is retained in mature sperm, providing a mechanistic link between paternal metabolic perturbation and the germline epigenome. Together, these findings demonstrate that folate availability shapes germline epigenome establishment through stage-specific metabolic and chromatin remodeling during spermatogenesis, revealing a metabolic basis for paternal environmental effects on the germline epigenome.

## Introduction

Environmental factors—including diet, lifestyle, and chemical exposures—affect the health of exposed individuals across the lifespan and, in some contexts, impact subsequent generations (1-3). Such acquired effects, excluding those resulting from genomic damage and mutation, are thought to be mediated by epigenetic memory, including DNA methylation, small RNAs, and histone modifications (4-7). While the Developmental Origins of Health and Disease (DOHaD) concept has highlighted the importance of the maternal environment (8, 9), growing evidence supports a complementary perspective—the Paternal Origins of Health and Disease (POHaD)(10)—in which paternal environmental exposures influence the health of offspring, potentially via sperm epigenome alterations (11). However, the molecular basis and developmental onset of paternal epigenomic perturbations during spermatogenesis remain poorly defined.

Paternal dietary imbalance has been reported as a major determinant of sperm DNA methylation, tRNA fragments, and histone methylation (12-16). Among nutritional factors, folate (vitamin B9) plays an essential role in the one-carbon metabolic pathway, which synthesizes S-adenosylmethionine (SAM), the universal methyl donor for methylation reactions (17). Consistently, folate availability has been shown to influence DNA and histone methylation in mouse sperm (18-20). Notably, a previous study demonstrated that dietary folate deficiency reduces sperm histone H3 lysine 4 trimethylation (H3K4me3) levels at enhancer regions of developmental genes and that these changes are partially transmitted to early embryos, leading to embryonic transcription errors and congenital abnormalities in the next generation (20). This provides a conceptual framework in which a portion of histone marks retained in sperm at regulatory regions of developmental genes may mediate paternal effects on offspring.

In mammalian sperm, 90–95% of histones are replaced by protamines (21), whereas a subset of nucleosomes frequently marked by H3K4me3 is retained at the developmental gene loci (22). This distinctive pattern of nucleosome retention in mature sperm is established throughout spermatogenesis (23, 24), a highly organized and continuous differentiation process. Spermatogonia first self-renew before retinoic acid–dependent commitment to meiotic entry. Spermatocytes then complete two meiotic divisions. Spermatids subsequently elongate into spermatozoa (25, 26). At each stage, locus-specific transcriptional activation and repression are coordinated by dynamic changes in chromatin landscape and histone modifications (27-29). Yet it remains unclear whether these chromatin and epigenetic dynamics depend on the metabolic state of testicular germ cells and at which stage nutritional perturbations alter the epigenome.

In this study, we show that dietary folate deficiency remodels epigenome establishment during spermatogenesis. Using unbiased metabolic profiling, we identify dynamic reprogramming of metabolic pathways across germ cell differentiation. We further demonstrate that metabolic perturbation reshapes multiple epigenetic layers, including chromatin accessibility, H3K4me3, and histone H3 lysine 27 trimethylation (H3K27me3). These alterations comprise genome-wide redistribution of H3K4me3, nuclear reorganization of H3K27me3, and partial persistence of H3K4me3 in sperm. Together, our findings reveal a metabolic basis for epigenome establishment in the male germline.

## Results

### Dynamics of metabolic pathways during mouse spermatogenesis

To identify stage-specific metabolic transitions that underlie epigenomic dynamics during spermatogenesis, we comprehensively profiled metabolites across spermatogenic differentiation. Testes were collected from neonatal and adult mice, and representative spermatogenic cell populations were isolated by fluorescence-activated cell sorting (FACS) for metabolomic analysis. Specifically, we analyzed undifferentiated spermatogonia (Undiff) and differentiated spermatogonia (Diff), both undergoing mitotic division; pachytene/diplotene spermatocytes (PD), corresponding to the middle of meiotic prophase I; and post-meiotic haploid round spermatids (RS) (Fig. 1A).

**Fig. 1.**
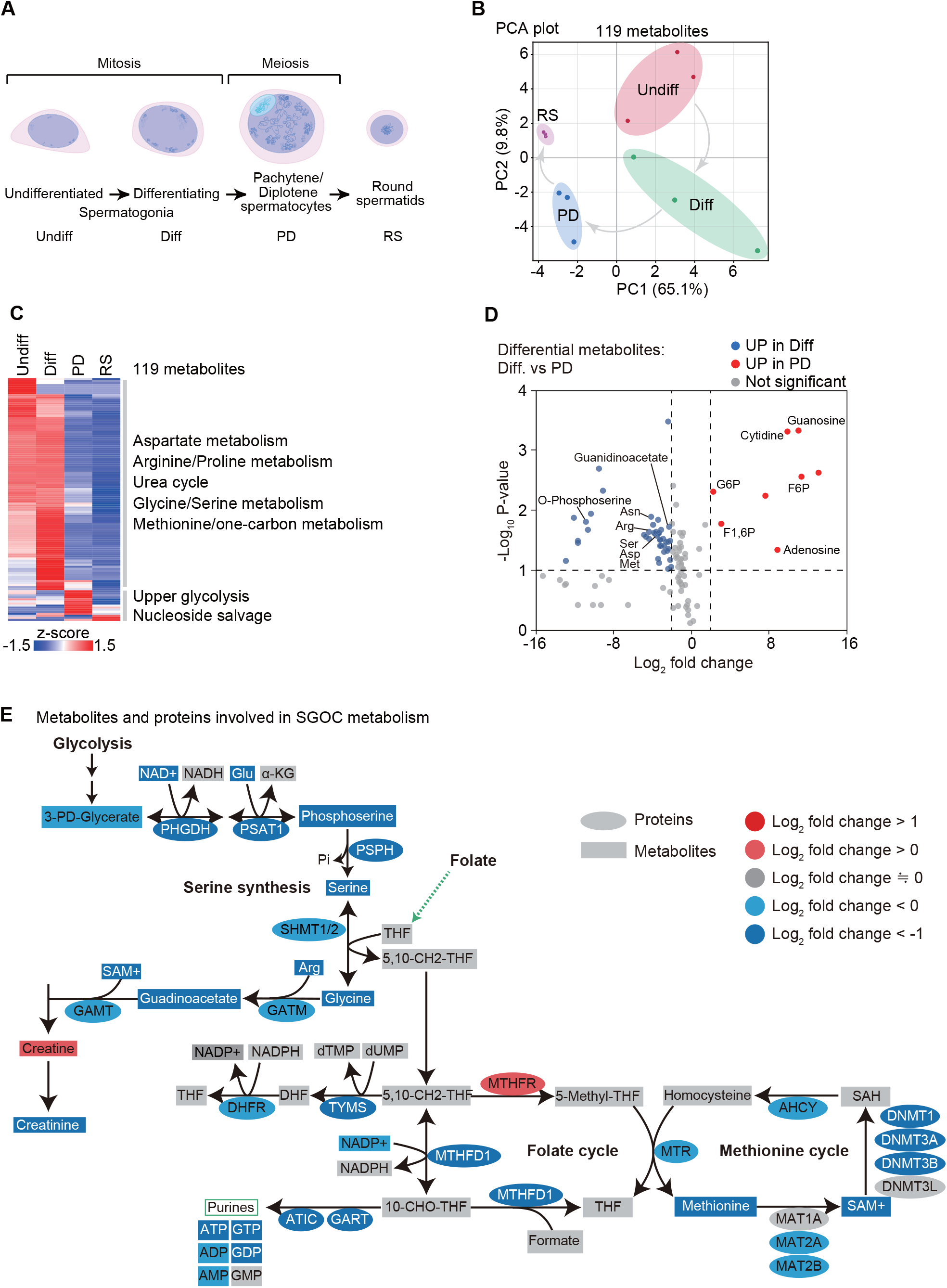
Stage-specific metabolic reprogramming during spermatogenesis. (A) Schematic of spermatogenesis and the four representative stages used for metabolomic analysis in this study. (B) Principal component analysis (PCA) of metabolomic profiles from four germ cell populations. Metabolomic measurements were performed in triplicate for each population. (C) Heatmap showing the relative abundance of 119 metabolites across the four germ cell populations. Metabolites detected in at least two of three measurements were retained for analysis. Metabolite levels were normalized by cell volume and visualized as row-wise Z-scores after hierarchical clustering. Clustering was performed using one minus Pearson correlation as the distance metric and average linkage. (D) Volcano plot comparing metabolite abundance between differentiating spermatogonia (Diff) and pachytene spermatocytes (PD). Each point represents a metabolite (x-axis, log_2_ fold change [PD/Diff]; y-axis, −log_10_ *P*-value from two-sided Student’s t-test). Metabolites enriched in Diff are shown in blue, those enriched in PD are shown in red, and metabolites without significant change are shown in gray. Significance thresholds were set at *P* < 0.1 and |log_2_ fold change| > 2. Selected metabolites belonging to serine–glycine–one-carbon (SGOC)-related and differentiation-associated metabolic pathways are indicated. (E) SGOC metabolic pathway illustrating metabolic and proteomic changes during the Diff-to-PD transition. Metabolites and enzymes detected in metabolomic and proteomic analyses were mapped onto the SGOC metabolic network. Colors represent log_2_ fold change between PD and Diff cells: dark red (>1), light red (>0), dark blue (<−1), light blue (<0), and gray indicates not detected.

In total, 140 metabolites were quantified, of which 119 metabolites detected in at least two biological replicates were retained for downstream analyses. Principal component analysis (PCA) revealed similar metabolic profiles between Undiff and Diff cells, whereas PD and RS formed distinct clusters (Fig. 1B), indicating a dramatic metabolic transition at the onset of meiosis. Consistent with this observation, hierarchical clustering of the 119 metabolites revealed pronounced shifts in metabolite abundance between mitotic spermatogonia and meiotic spermatocytes (Fig. 1C). Notably, the majority of metabolites displayed reduced abundance during the Diff-to-PD transition. Metabolites involved in serine–glycine–one-carbon (SGOC) metabolism showed particularly strong reductions. Because SGOC metabolism lies downstream of glycolysis and links cellular metabolic state to epigenetic regulation (30, 31), these observations suggest that metabolic changes during spermatogenesis may be coupled to shifts in epigenetic regulation (28, 32). Differential analysis between Diff and PD cells further confirmed that a large fraction of metabolites decreased in abundance during this transition (Fig. 1D), suggesting widespread metabolic downregulation at the pachytene stage. Together, these results indicate that metabolic programs, similar to transcriptional programs, undergo coordinated and stage-specific remodeling during the mitotic, meiotic, and haploid phases of spermatogenesis.

To identify metabolic pathways underlying these changes, we performed enrichment and pathway analyses on metabolites reduced during meiotic prophase I. These analyses revealed significant alterations in glycolysis and glycine, serine, and threonine metabolism during the transition from mitotic spermatogonia to meiotic spermatocytes (Fig. S1), indicating reprogramming of energy-producing pathways at meiotic onset. Among these pathways, glycine, serine, and threonine metabolism are of particular interest because the SGOC pathway contributes to the synthesis of SAM, a universal methyl donor for DNA and histone methylation. We therefore focused subsequent analyses on SGOC metabolism.

Integrated analyses of metabolomic, proteomic, and transcriptomic datasets revealed coordinated reductions in metabolites and enzymes across multiple nodes of the SGOC pathway during meiosis (Fig. 1E). These changes spanned the glycolysis-linked serine synthesis branch, the folate cycle, and the methionine cycle, including decreases in serine and glycine metabolism, folate-cycle intermediates, and components involved in SAM production. Together, these alterations indicate a global attenuation of SGOC metabolic flux in meiotic spermatocytes. Collectively, these results indicate that SGOC metabolism is active during the mitotic phase of spermatogenesis and is selectively downregulated in meiotic spermatocytes.

### Dietary folate deficiency model and spermatogenesis

Given that SGOC metabolism was substantially reduced at the transition from mitotic spermatogonia to meiotic spermatocytes, we hypothesized that folate availability, a key determinant of SGOC metabolism, may influence spermatogenic differentiation and epigenome. To test this, we examined the effects of folate deficiency on spermatogenesis. In our folate-deficient mouse model, male mice at 3 weeks of age, immediately after weaning, were fed either a folate-deficient (FD) diet (18, 20) or a folate-sufficient (FS) control diet (33) for 6 weeks, a period covering spermatogenic differentiation (34, 35) and enabling assessment of spermatogenesis under folate-deficient conditions (Fig. S2A and B). Body weight did not differ between FD and FS mice throughout the feeding period (Fig. S2C). Following dietary treatment, males were mated with FS-fed females, after which testes and caudal epididymides were collected for analysis. No significant differences were observed between the two groups in testis weight, sperm count, or litter size (Fig. S2D–F). In addition, immunohistochemical analyses revealed no changes in the proportions of germ cells at distinct stages of the seminiferous epithelial cycle (stages I–III, IV–VII, and VIII–IX) (36) under folate-deficient conditions (Fig. S2G and H). Together, these results demonstrate that dietary folate deficiency initiated after weaning does not measurably impair spermatogenic progression or male fertility, providing a suitable experimental framework for focusing on epigenomic profiles in the absence of overt reproductive defects.

### Folate deficiency alters genome-wide chromatin accessibility during spermatogenesis

To determine whether folate deficiency perturbs the epigenomic landscape during spermatogenesis, we performed ATAC-seq to profile genome-wide chromatin accessibility, a functional readout of gene regulatory element activity (37, 38). Chromatin accessibility was profiled across representative stages of spermatogenesis, including Undiff, Diff, leptotene/zygotene spermatocytes (LZ), PD, and RS (Fig. S3A and B). Only modest changes were detected from Undiff through LZ (266 upregulated and 89 downregulated differentially accessible regions (DARs) in Undiff; 765 upregulated and 972 downregulated DARs in Diff; and 39 upregulated and 4 downregulated DARs in LZ). In contrast, substantial alterations were observed in PD and RS (5,105 upregulated and 4,114 downregulated DARs in PD; 2,951 upregulated and 5,600 downregulated DARs in RS) (Fig. 2A), indicating that folate deficiency alters genome-wide chromatin accessibility beginning in meiotic spermatocytes.

**Fig. 2.**
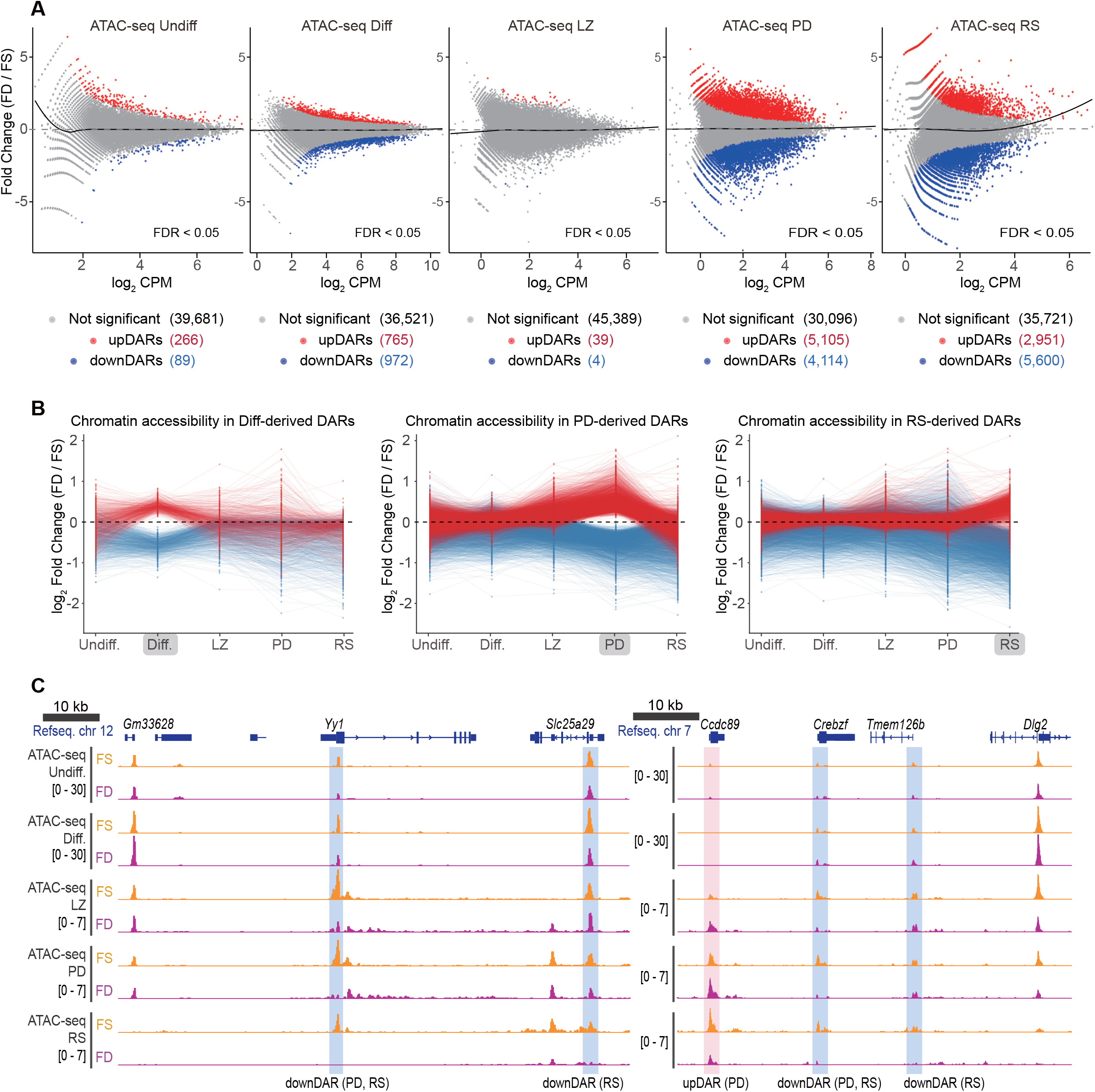
Chromatin accessibility changes during spermatogenesis under folate deficiency. (A) MA plots showing differential chromatin accessibility between folate-sufficient (FS) and folate-deficient (FD) cells at five differentiation stages (Undiff., Diff., LZ, PD, and RS). Each dot represents an ATAC-seq peak region at each stage. The x-axis indicates average accessibility (log2 CPM), and the y-axis indicates the log2 fold change (FD vs. FS); the dashed line indicates zero-fold change. Regions with FDR < 0.05 were defined as differentially accessible regions (DARs), with increased accessibility in FD (upDARs; red) and decreased accessibility in FD (downDARs; blue). Not significant regions are shown in gray. The black curve indicates the global trend across all regions. (B) Trajectories of ATAC-seq log2 fold changes (FD vs. FS) at DARs identified in Diff., PD, and RS. Each line connecting dots represents an individual upDAR (red) or downDAR (blue) based on accessibility changes across the three stages. The y-axis shows log2(FD/FS), and the dashed line indicates no change. (C) Genome browser tracks showing ATAC-seq signals in FS (yellow) and FD (pink) cells across five differentiation stages. Signals are displayed as CPM-normalized coverage. Regions highlighted in red and blue correspond to upDARs and downDARs, respectively. RefSeq gene models are shown at the top, and the black bar indicates genomic scale.

We next examined whether these accessibility changes were transient or sustained across differentiation. In Diff, which showed modest changes, and in PD and RS, which exhibited the greatest numbers of DARs, increases in chromatin accessibility were largely stage-specific and transient (red lines in Fig. 2B). In contrast, decreases in chromatin accessibility tended to persist in later stages. Reductions detected in Diff remained decreased in subsequent PD and RS stages, and prominent reductions in PD and RS were broadly maintained across stages (blue lines in Fig. 2B). Representative genomic track views confirmed that regions with decreased accessibility in PD persisted into RS (Fig. 2C). Collectively, these findings demonstrate that dietary folate deficiency induces widespread remodeling of chromatin accessibility during spermatogenesis, with the onset of prominent changes at pachytene/diplotene stage and a distinct pattern of transient gains but persistent losses.

### Folate deficiency induces cell-type–specific remodeling of chromatin accessibility

Given the contrasting pattern of up- and downregulated DARs detected in PD and RS, we next explored the gene regulatory programs associated with these accessibility changes. Each set of DARs was assigned to putative cis-regulatory elements of nearby genes using GREAT analysis (39). Upregulated and downregulated DARs in PD and RS exhibited distinct genomic distributions. DownDARs were preferentially localized near promoter–transcription start site (TSS), whereas upDARs were more frequently detected in intergenic regions (Fig. 3A and Fig. S4A).

**Fig. 3.**
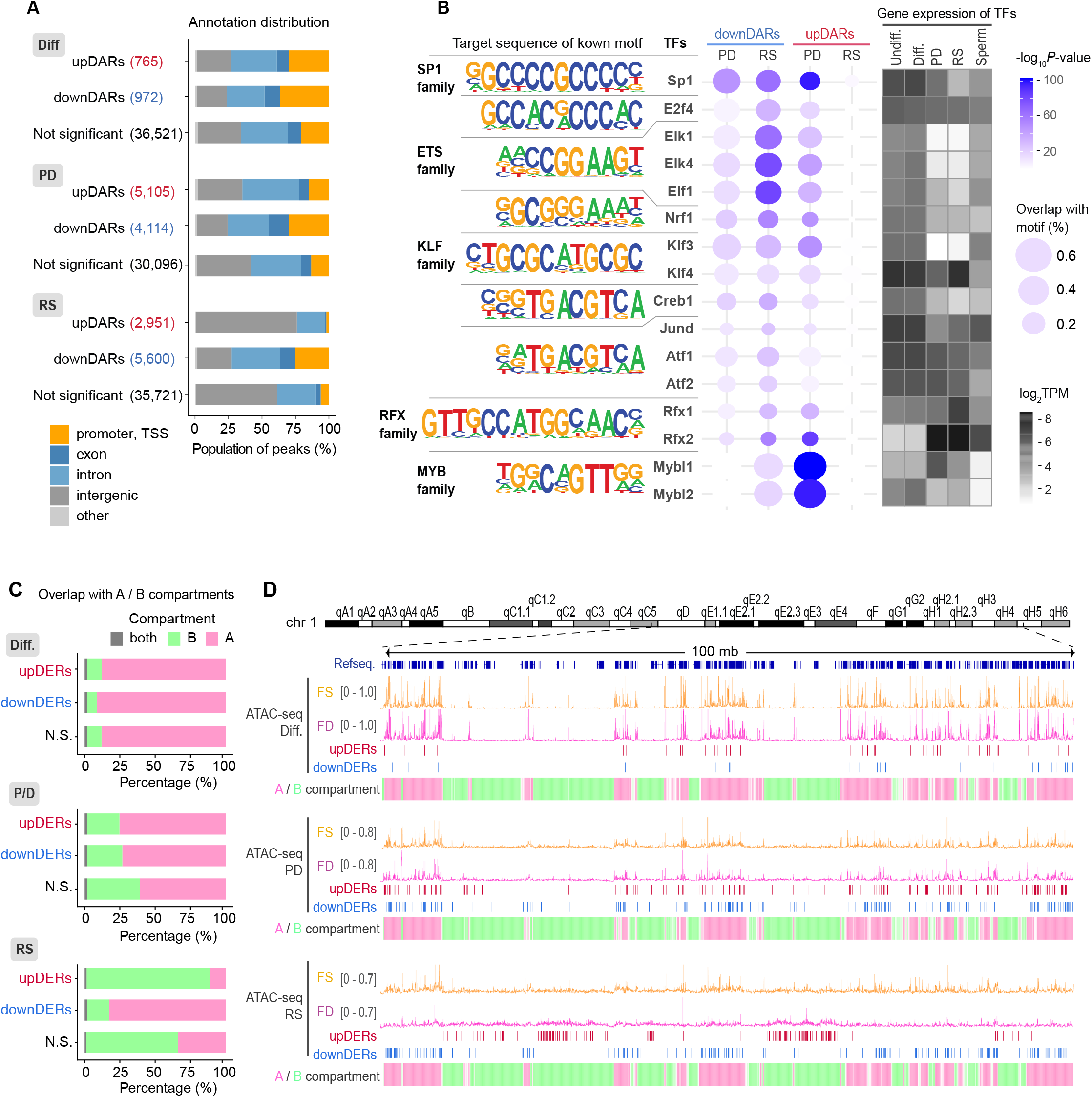
Genomic features and transcription factors of DARs. (A) Distribution of genomic annotations for upDARs, downDARs, and not significant regions in Diff., PD, and RS. Peaks were categorized into promoter–TSS, exon, intron, intergenic, and other regions, based on RefSeq gene annotation. Bars represent the percentage of regions within each category. (B) Putative transcription factors (TFs) enriched in DARs in PD and RS are shown as a balloon plot. DNA sequence logos represent TF-binding motifs identified using the HOMER package. Bubble size indicates the percentage of target sequences containing the motif, and color represents the –log10 P-value. Germ cell–expressed TFs among the top-enriched motifs are highlighted and shown with their expression levels (log2 TPM) across spermatogenic stages in the accompanying heatmap. (C) Chromatin compartment distribution of DARs based on differential enrichment between FS and FD. DARs in Diff, PD, and RS were intersected with A/B chromatin compartments defined from Hi-C data in round spermatids (50-kb resolution). The proportion of peaks overlapping A or B compartments was calculated for each category. Regions overlapping multiple compartments were labeled as “both.” Bars indicate the percentage of peaks within each compartment. (D) Genome browser tracks showing ATAC-seq signals in FS (yellow) and FD (pink) samples across Diff, PD, and RS stages with the locations of DARs. ATAC-seq signals are displayed as CPM-normalized coverage. The Hi-C–derived PCA1 track (bedGraph) is shown below, where positive PCA1 values (pink) correspond to A compartments and negative PCA1 values (green) correspond to B compartments. Chromosomal coordinates and RefSeq gene models are shown at the top, and the black bar indicates the genomic scale.

Genes associated with PD upDARs were enriched for cilium movement and microtubule-based processes, suggesting a potential involvement of axonemal and flagellar gene programs required for sperm motility (Fig. S4B). In contrast, RS upDAR–associated genes were enriched for plasma membrane adhesion pathways. Conversely, genes linked to downDARs across Diff, PD, and RS were consistently enriched for RNA-related processes, including RNA stability and RNA metabolic pathways, suggesting coordinated repression of RNA regulatory programs.

To further explore the regulatory mechanisms underlying these accessibility changes, we performed motif enrichment analysis (40) within DARs. Motifs for SP1, ETS, and KLF family transcription factors (TFs) were enriched across multiple DAR sets. Notably, binding motifs for MYB and Regulatory Factor X (RFX) family TFs were specifically enriched in PD upDARs (Fig. 3B), suggesting a potential role for meiotic transcriptional regulators in folate-responsive chromatin remodeling. In Diff downDARs, DNMT1- and BORIS/CTCFL-associated sites were enriched, indicating that the impact of folate deficiency on chromatin accessibility is cell-type specific (Fig. S4B).

Because RS upDARs lacked enriched TF motifs, TF binding alone may not account for these accessibility changes, and we therefore examined whether higher-order chromatin organization might contribute to their regulation. Reanalysis of previously published Hi-C data from wild-type spermatogenic cells (41) revealed that more than 80% of RS upDARs were located within B compartments, whereas the majority of other DARs resided in A compartments (Fig. 4C). Track-view visualization further showed that RS upDARs were frequently positioned near the centers of B compartments (Fig. 4D). Because B compartments correspond to gene-poor and transcriptionally inactive chromatin domains (42), this preferential localization in B compartments suggests that folate deficiency may also affect repressive chromatin environments in RS. Furthermore, considering the absence of enriched TF motifs (Fig. 3B), RS upDARs may reflect chromatin architecture–dependent mechanisms rather than canonical TF–driven regulation. Together, these findings indicate that dietary folate deficiency induces stage-specific remodeling of chromatin accessibility during spermatogenesis, involving distinct genomic localization patterns, TF programs, and chromatin compartment contexts.

**Fig. 4.**
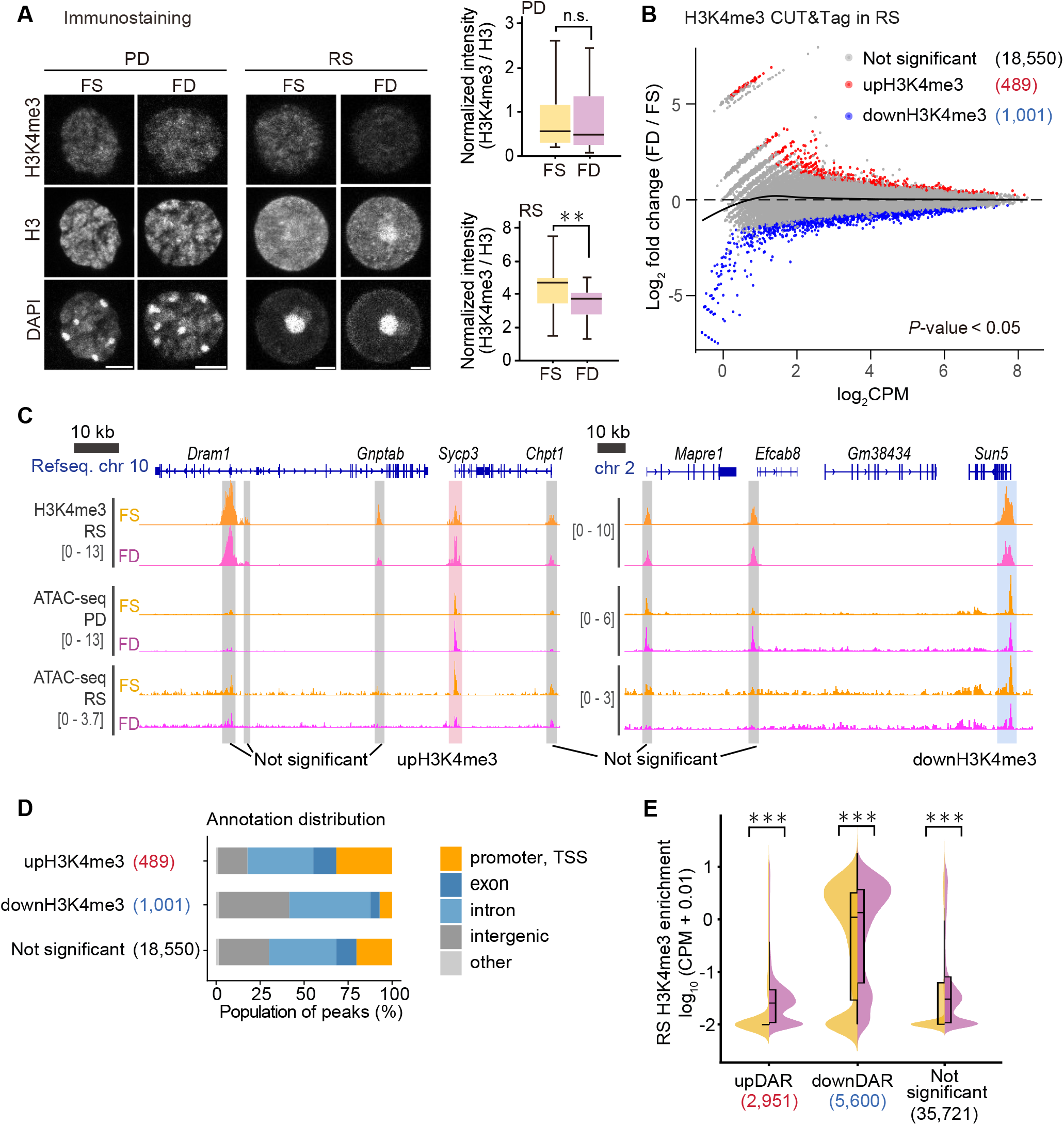
H3K4me3 remodeling in post-meiotic cells. (A) Immunocytochemical analysis of H3K4me3 in PD and RS prepared as single-cell suspensions from FS and FD testes. Cells were stained for H3K4me3, H3, and DAPI, and imaged by confocal microscopy. PD and RS were distinguished based on characteristic nuclear morphology. Representative grayscale images are shown for each marker. Approximately 50 cells per group with comparable H3 signal intensity were selected for analysis to minimize potential signal variability derived from focal depth and total histone abundance. Regions of interest (ROIs) were defined within the nucleus, and background-subtracted H3K4me3 signal intensity was normalized to H3 intensity for each cell (H3K4me3/H3 ratio). Box plots show the distribution of normalized H3K4me3 intensity (H3K4me3/H3) in PD and RS under FS and FD conditions. Statistical significance was assessed using a two-sided Student’s *t*-test. No significant difference was observed in PD (P = 0.98, n.s.), whereas a significant difference was detected in RS (P = 0.0002, ***). (B) MA plot showing differential H3K4me3 enrichment between FS and FD cells in RS. Each dot represents a H3K4me3 peak region defined from the union peak set. The x-axis indicates average H3K4me3 signal intensity (log2 CPM), and the y-axis indicates the log2 fold change (FD vs. FS); the dashed line indicates zero-fold change. Regions with *P* < 0.05 were defined as differentially enriched regions, with increased H3K4me3 enrichment in FD (upH3K4me3; red) and decreased enrichment in FD (downH3K4me3; blue). Not significant regions are shown in gray. The black curve indicates the global trend across all regions. (C) Genome browser tracks showing H3K4me3 CUT&Tag signals in RS and ATAC-seq signals in PD and RS, colored in FS (yellow) and FD (pink) cells. Signals are displayed as CPM-normalized coverage. Regions highlighted in red and blue correspond to upH3K4me3 and downH3K4me3, respectively. RefSeq gene models are shown at the top, and the black bar indicates genomic scale. (D) Distribution of genomic annotations for upH3K4me3, downH3K4me3, and not significant regions in RS. Peaks were categorized into promoter–TSS, exon, intron, intergenic, and other regions based on RefSeq gene annotation. Bars represent the percentage of regions within each category. (E) Violin plots showing the distribution of H3K4me3 enrichment in RS across ATAC-defined DAR classes. For each DAR class, the average H3K4me3 CUT&Tag signal was quantified and compared between FS and FD RS. Signal intensity is shown as log□□-transformed normalized coverage with a pseudocount added (log□□[signal + 0.01]) to avoid zero values. Violin plots represent distribution density, with overlaid boxplots indicating the median and interquartile range. Statistical significance between FS and FD was assessed using a two-sided Wilcoxon rank-sum test and is simply shown (*P < 0.05, **P < 0.01, ***P < 0.001, n.s., not significant).

### Folate deficiency disturbs genome-wide H3K4me3 independently of chromatin accessibility in post-meiotic cells

To determine whether folate deficiency alters histone methylation in addition to chromatin accessibility, we examined H3K4me3, a chromatin mark associated with active regulatory regions (43) and linked to folate-dependent one-carbon metabolism. Because H3K4 trimethylation is catalyzed by SET1/COMPASS methyltransferases that require SAM as a methyl donor (44, 45), we hypothesized that folate deficiency may perturb H3K4me3 levels during spermatogenesis. Immunofluorescence analysis revealed a significant reduction in global H3K4me3 signal in RS under FD conditions, whereas PD showed no apparent change (Fig. 4A). To further examine these changes at the genomic level, we performed H3K4me3 CUT&Tag (46) in RS.

Genome-wide analysis revealed that H3K4me3 was altered in both directions under FD conditions: while many peaks decreased (downH3K4me3, n = 1,001), a subset of peaks increased (upH3K4me3, n = 489) (Fig. 4B and C; Fig. S5A). UpH3K4me3 peaks were predominantly located in intergenic regions, whereas downH3K4me3 peaks were enriched at promoter–TSS (Fig. 4D), consistent with previous observations of H3K4me3 alterations in epididymal sperm from FD mice (20). Notably, the genomic distribution of H3K4me3 alterations did not directly mirror the chromatin accessibility changes detected by ATAC-seq. To evaluate the relationship between these two epigenomic layers, we compared CUT&Tag and ATAC-seq signals genome-wide. H3K4me3 enrichment was increased in FD relative to FS mice, particularly at regions corresponding to upDARs (Fig. 4E). In contrast, H3K4me3 levels within downDARs were only modestly altered by folate deficiency. DownDARs were frequently located within regions exhibiting high basal H3K4me3 enrichment and were associated with specific TF motifs (Fig. 3C and Fig. S5B). Collectively, these findings demonstrate that dietary folate deficiency induces region-specific gains and losses of H3K4me3 in post-meiotic germ cells and that these changes are not simply a consequence of altered chromatin accessibility, indicating partially independent layers of epigenomic remodeling. These bidirectional changes suggest that folate deficiency does not uniformly reduce histone methylation but instead reshapes locus-specific H3K4me3 landscapes.

### Folate deficiency alters nuclear organization of H3K27me3 without widespread redistribution of bivalent domain

During spermatogenesis, H3K4me3 frequently colocalizes with H3K27me3 at developmental gene loci (28, 47, 48), forming bivalent domains that antagonistically regulate transcriptional activity and chromatin accessibility (29). Because H3K4me3 levels were altered in RS under FD conditions, we next examined whether H3K27me3 was similarly affected. Immunofluorescence analysis revealed ectopic H3K27me3-positive nuclear aggregates in RS from FD testes (Fig. 5A). These aggregates were distinct from chromocenters (CC) (49) and post-meiotic sex chromosome (PMSC) (50) structures but resembled facultative heterochromatin foci previously reported in RS lacking the germline Polycomb component SCML2, a germline-specific regulator of H3K27me3 (51).

**Fig. 5.**
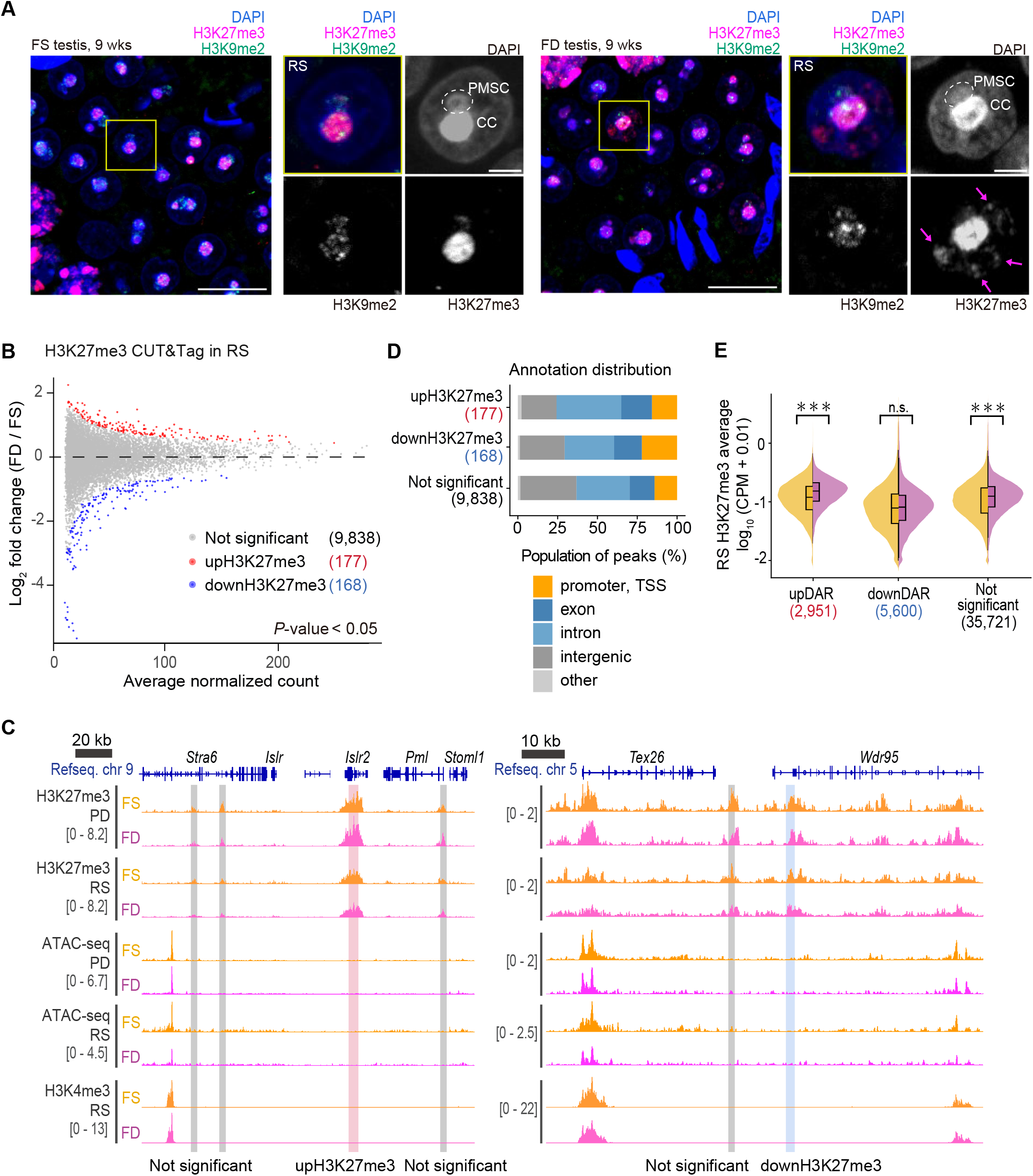
Nuclear reorganization of H3K27me3 in FD spermatids. (A) Immunohistochemical images of testis sections from 9-week-old FS and FD mice show the nuclear distribution of H3K27me3. Representative seminiferous tubules at stages I–III are shown in the left panels (scale bar, 10 μm), highlighting the eary-RS cell population. Higher-magnification views of representative cells (yellow-boxed areas) are displayed on the right (scale bar, 2 μm). Images are presented as single-channel grayscale and merged views. H3K27me3 is shown in magenta, H3K9me2 in green, and DAPI in blue. Magenta arrows indicate ectopic H3K27me3 patches specifically observed in early RS of FD testes. (B) MA plot showing differential H3K27me3 enrichment between FS and FD cells in RS. Each dot represents a broad H3K27me3 peak region derived from the consensus peak set. The x-axis indicates the average of normalized raw counts (baseMean) calculated by DESeq2, and the y-axis shows the log2 fold change (FD vs. FS). The dashed horizontal line denotes no change (log2FC = 0). Regions with *P* < 0.05 were defined as differentially enriched, with increased H3K27me3 levels in FD (upH3K27me3; red) and decreased levels in FD (downH3K27me3; blue), while non-significant regions are shown in gray. Differential analysis was performed using DESeq2 on raw count matrices derived from broad peak regions, incorporating size-factor normalization and surrogate variable adjustment to account for latent batch effects. (C) Genome browser tracks showing H3K27me3 CUT&Tag signals in PD and RS, H3K4me3 CUT&Tag signals in RS, and ATAC-seq signals in PD and RS. Tracks are colored by condition: FS (yellow) and FD (pink). Signals are displayed as CPM-normalized coverage. Regions highlighted in red and blue correspond to upH3K27me3 and downH3K27me3, respectively. RefSeq gene models are shown at the top, and the black bar indicates genomic scale. (D) Distribution of genomic annotations for upH3K27me3, downH3K27me3, and not significant regions in RS. Peaks were categorized into promoter–TSS, exon, intron, intergenic, and other regions based on RefSeq gene annotation. Bars represent the percentage of regions within each category, consistent with the broad genomic distribution characteristic of H3K27me3 domains. (E) Violin plots showing the distribution of H3K27me3 enrichment in RS across ATAC-defined DAR classes. For each DAR class, the average H3K27me3 CUT&Tag signal was quantified and compared between FS and FD RS. Signal intensity is shown as log□□-transformed normalized coverage with a pseudocount added (log□□[signal + 0.01]) to avoid zero values. Violin plots represent distribution density, with overlaid boxplots indicating the median and interquartile range. Statistical significance between FS and FD was assessed using a two-sided Wilcoxon rank-sum test and is indicated using standard notation (*P < 0.05, **P < 0.01, ***P < 0.001, n.s., not significant).

In *Scml2*-knockout testes, global H3K27me3 levels are reduced in PD and RS (28). Based on these observations, we performed H3K27me3 CUT&Tag in PD and RS to define genome-wide changes (Fig. S6A and B). Unexpectedly, genome-wide profiling revealed only limited alterations in H3K27me3 occupancy under FD conditions. In RS, 177 peaks showed increased H3K27me3 and 168 showed decreased H3K27me3 at a threshold of P < 0.05 (Fig. 5B). Similarly, PD exhibited only modest changes (116 upH3K27me3 and 117 downH3K27me3 peaks; Fig. S6C). Annotation analysis showed comparable genomic distributions for up- and downregulated H3K27me3 peaks in both PD and RS (Fig. 5C and D; Fig. S6D), indicating no major redistribution of H3K27me3 across genomic features.

We further examined H3K4me3 and H3K27me3 enrichment at TSSs of genes known to form bivalent domains during spermatogenesis (28). Under FD conditions, H3K4me3 levels were modestly increased at these loci, whereas H3K27me3 levels remained largely unchanged. In contrast, chromatin accessibility at the same loci was markedly reduced in FD testes (Fig. S6E and F). Collectively, these results indicate that dietary folate deficiency alters the nuclear organization of H3K27me3 and substantially affects chromatin accessibility in RS, despite causing only minimal changes in genome-wide H3K27me3 occupancy in meiotic and post-meiotic germ cells.

### FD-induced H3K4me3 alterations in post-meiotic cells are partially retained in sperm

Finally, to determine whether the epigenetic changes observed in testis are retained in mature sperm and may contribute to paternal epigenetic inheritance, we compared our H3K4me3 CUT&Tag data with previously published H3K4me3 ChIP-seq data from epididymal sperm of FD mice (20). Using the published dataset, we defined sperm upH3K4me3 (n = 534) and downH3K4me3 (n = 869) peaks at a threshold of P < 0.05 (Fig. 6A). Consistent with previous reports (20), sperm upH3K4me3 peaks were predominantly enriched at promoter regions, whereas downH3K4me3 peaks were more frequently localized to intergenic regions (Fig. 6B).

**Fig. 6.**
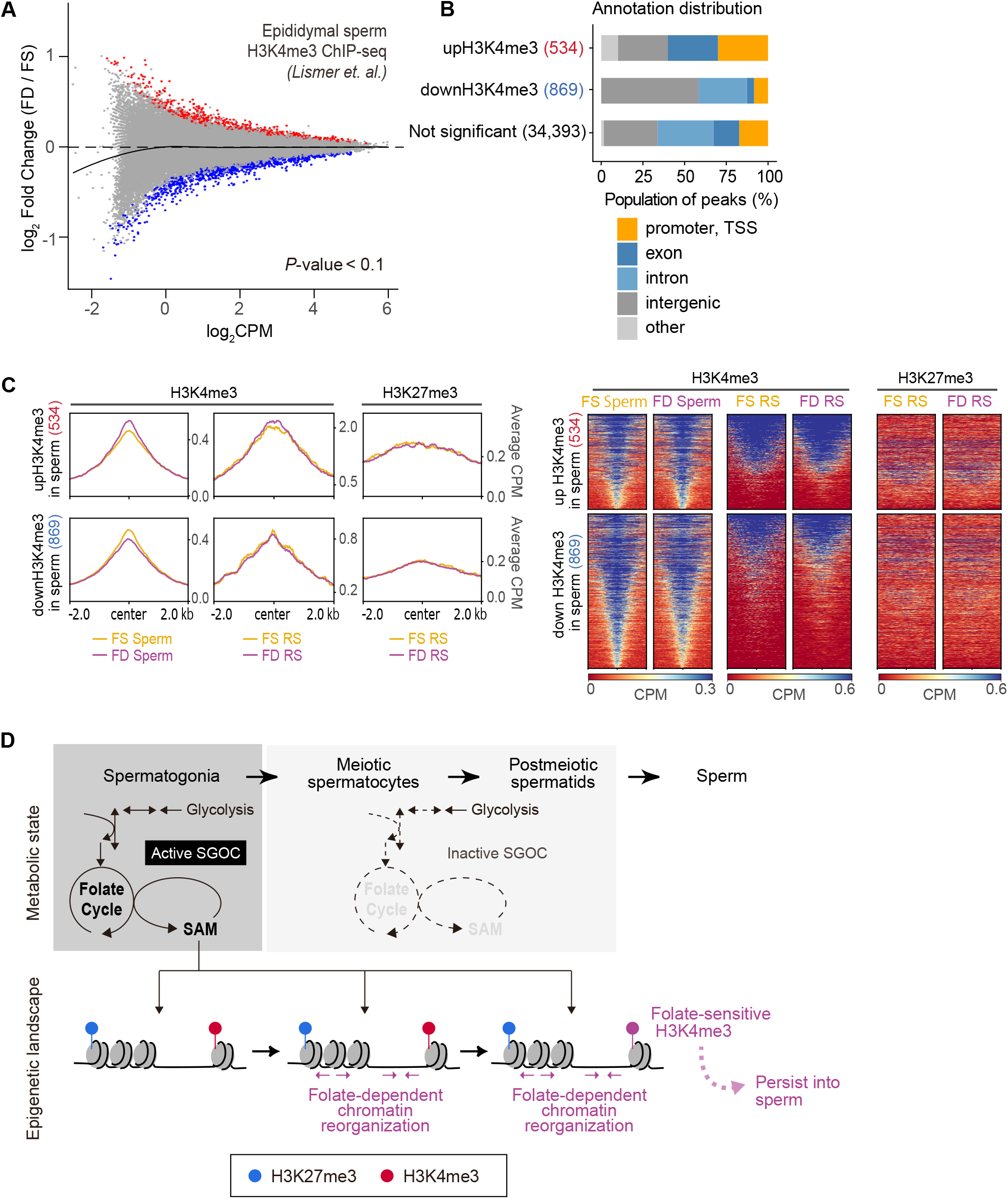
Partial retention of FD-induced H3K4me3 changes in sperm. (A) MA plot illustrating differential H3K4me3 enrichment between FS and FD conditions based on reanalysis of previously published sperm H3K4me3 ChIP-seq datasets. Each dot represents a H3K4me3 peak defined from the consensus peak set derived from sperm. The x-axis indicates the average H3K4me3 signal intensity (log2 CPM), and the y-axis shows the log2 fold change (FD vs. FS), with the dashed line marking no change (log2FC = 0). Regions with *P* < 0.1 were considered differentially enriched, including loci with increased H3K4me3 levels in FD (upH3K4me3 in sperm; red) and decreased levels in FD (downH3K4me3 in sperm; blue), while not significant regions are shown in gray. The black curve represents the overall trend across all sperm H3K4me3 peaks. (B) Genomic annotation distribution of upH3K4me3, downH3K4me3, and non-significant regions identified in sperm. Peaks were classified into promoter–TSS, exon, intron, intergenic, and other categories according to RefSeq gene annotation. Bars indicate the proportion of peaks within each genomic category. (C) Average tag density profiles of H3K4me3 and H3K27me3 CUT&Tag signals in RS centered on sperm H3K4me3 differential regions (upH3K4me3 and downH3K4me3 in sperm). Signal intensity was calculated across ±2 kb from the peak center using a reference-point approach. Profiles are shown separately for two regions, and the y-axis represents average signal intensity. Heatmap representation of H3K4me3 and H3K27me3 CUT&Tag signals in RS across sperm H3K4me3 differential regions (upH3K4me3 and downH3K4me3 in sperm), aligned at the peak center and displayed over a ±2 kb window. Signal intensities are shown as normalized coverage, with the color scale capped at a maximum value. Each row represents an individual peak region, grouped according to up and down classification. (D) Conceptual model of folate-dependent epigenome remodeling during spermatogenesis. Metabolomic analysis reveals stage-specific metabolic remodeling, including reduced SGOC metabolism upon entry into meiosis, potentially limiting SAM availability. Under folate-deficient conditions, this metabolic transition coincides with extensive reorganization of chromatin accessibility and histone modifications, particularly in meiotic and post-meiotic cells. The active histone mark H3K4me3 undergoes stage-specific redistribution in early spermatids. A subset of these H3K4me3 alterations is retained in mature sperm, suggesting a potential mechanism linking paternal folate status to the germline epigenome.

To assess whether RS alterations are retained in sperm, we examined H3K4me3 and H3K27me3 enrichment at sperm H3K4me3-altered regions. At sperm upH3K4me3 loci, RS H3K4me3 exhibited a concordant increasing trend under folate deficiency, whereas RS H3K27me3 levels remained largely unchanged (Fig. 6C and Fig. S7). Similarly, sperm downH3K4me3 regions showed only modest differences in RS H3K4me3 levels but not in RS H3K27me3 levels. Taken together, these results indicate that a subset of FD-induced H3K4me3 alterations in post-meiotic RS is retained in mature sperm, particularly at regions exhibiting increased H3K4me3, supporting the model that selective histone methylation changes established during spermatogenesis can persist into mature sperm under folate-deficient conditions (Fig. 6D).

## Discussion

Intergenerational and transgenerational effects of parental nutrition are increasingly recognized in epidemiological studies and represent a significant risk factor for congenital disorders (52, 53). In a mouse model of paternal folate deficiency initiated at weaning, dietary folate deprivation induces epigenomic alterations in sperm, leading to aberrant gene expression in early embryos and craniofacial malformations in offspring (20). However, the mechanistic link between dietary folate deficiency and sperm epigenomic remodeling has remained unclear. In this study, we sought to define how folate availability is translated into stage-specific metabolic and epigenomic alterations during spermatogenesis.

Our metabolomic analyses reveal that metabolic landscapes undergo coordinated and stage-specific remodeling during spermatogenic differentiation, with a pronounced transition from mitotic spermatogonia to meiotic spermatocytes. Undifferentiated and differentiated spermatogonia share highly similar metabolic profiles, whereas meiotic and post-meiotic cells exhibit distinct metabolic states, indicating structured metabolic reprogramming aligned with the mitotic, meiotic, and haploid phases of germ cell differentiation. This transition is characterized by a global reduction in metabolite abundance and coordinated changes in energy-producing pathways, including glycolysis and glycine, serine, and threonine metabolism. Notably, SGOC metabolism, which links cellular metabolic states to epigenetic regulation through SAM synthesis, is selectively active during the mitotic phase and globally downregulated during the transition from mitotic spermatogonia to meiotic spermatocytes. Consistent with this organization, folate levels are enriched in spermatogonia and decline during meiotic and post-meiotic stages, revealing a developmentally programmed reorganization of one-carbon metabolism during spermatogenesis.

Despite enrichment of SGOC metabolites in spermatogonia, folate deficiency–induced epigenomic alterations become evident only after late meiosis, particularly in pachytene/diplotene spermatocytes and round spermatids. This temporal uncoupling suggests a model of metabolic priming, in which metabolites accumulated during the mitotic phase support chromatin regulation during later stages of germ cell differentiation. Notably, the observed tendency for folate deficiency– induced losses in chromatin accessibility was persistent from spermatogonia to RS, supporting the concept that metabolic perturbation affects epigenetic priming (54) across differentiation stages of spermatogenesis.

Chromatin accessibility profiling identifies multiple patterns of folate-sensitive remodeling. Accessibility gains in meiotic spermatocytes are largely transient and enriched at intergenic regions, including loci associated with sperm motility– related gene networks. Such changes may reflect compensatory epigenomic responses that preserve spermatogenic progression, consistent with the absence of overt fertility defects in this model. Accessibility gains in round spermatids are also transient and predominantly intergenic but occur near developmental regulatory loci, suggesting that germline epigenomic programming at this stage may preferentially affect developmental gene networks. In contrast, accessibility losses are persistent across differentiation stages and enriched at promoters of genes involved in RNA processing and RNA stability. These coordinated changes raise the possibility that folate deficiency may influence post-transcriptional regulatory pathways, including small RNA dynamics that have been implicated in paternal inheritance (55).

In addition to chromatin accessibility remodeling, we observe redistribution of the active histone mark H3K4me3 in round spermatids as well as altered nuclear organization of H3K27me3. Although genome-wide H3K27me3 occupancy remains largely stable, folate deficiency induces ectopic H3K27me3-positive nuclear aggregates resembling facultative heterochromatin foci described in Polycomb-deficient germ cells (51). Because these aggregates are not accompanied by widespread peak-level changes, they may reflect higher-order chromatin structural alterations rather than canonical redistribution of Polycomb targets. Moreover, similar nuclear puncta and compartment-dependent remodeling have been reported in aging-associated chromatin reorganization (56), suggesting that folate deficiency may induce a premature or stress-associated chromatin compaction phenotype in post-meiotic cells. Similarly, perturbation of the counterbalancing relationship between H3K4me3 and H3K27me3 at developmental loci has been proposed as a driver of heritable epigenetic instability (57). These findings suggest that folate deficiency perturbs chromatin organization at multiple levels, including both regulatory element accessibility and histone modification landscapes.

Notably, genome-wide H3K4me3 alterations observed in round spermatids are partially retained in mature sperm. However, chromatin accessibility changes detected in testis do not uniformly correspond to H3K4me3 alterations in sperm, indicating that multiple epigenomic layers may contribute independently to germline inheritance. Previous studies have shown that environmental perturbations can influence sperm small RNA composition and thereby mediate intergenerational phenotypes (12, 15, 16), raising the possibility that distinct epigenetic carriers operate in parallel.

Together, this study provides an integrated framework linking dietary folate deficiency to epigenomic regulation through stage-specific metabolic programming during spermatogenesis. By defining when and how folate-dependent one-carbon metabolism interfaces with chromatin architecture, our findings advance mechanistic understanding of how paternal nutrition shapes the germline epigenome and potentially influences offspring development.

## Materials and methods

### Mice

Mice were housed under a controlled 12 h light/12 h dark cycle at 21°C and provided with food and water *ad libitum*. After the dietary intervention, all mice were fed a normal diet (MR standard; Sankyo Labo Service Corporation). All animal procedures were approved by the guidelines of the Institutional Animal Care and Use Committee at Tokyo University of Science.

### Dietary treatment

Male *C57BL/6NCrslc* mice (Sankyo Labo Service Corporation) were randomly allocated to two dietary groups. The folate-sufficient (FS) group was fed a control diet (AIN-93G; CLEA Japan) containing 2 mg/kg folic acid (33), whereas the folate-deficient (FD) group was fed a folic-acid-free AIN-93G (CLEA Japan) containing 0 mg/kg folic acid (S. Fig. 2B). Dietary exposures in male mice were started at 3 weeks of age and lasted for 6 weeks. During the dietary phase, body weight of each mouse was monitored weekly (S. Fig. 2A and 2C).

### Isolation of testicular germ cells

For metabolomic analysis, spermatogonia were isolated from testes of mice at postnatal day 7 (P7), while pachytene spermatocytes and round spermatids were isolated from adult testes (8–12 weeks old). For next-generation sequencing (NGS) analysis, male germ cells at each developmental stage were isolated from adult testes (9 weeks old) after dietary exposure. Firstly, the tunica albuginea membrane was removed, and the seminiferous tubules were collected in Dulbecco’s Modified Eagle Medium (DMEM; WAKO) and minced several times. Tubules were washed three times with ice-cold DMEM to remove testicular sperm, followed by digestion in DMEM containing 1 mg/mL collagenase type IV (Worthington) and 20 µg/mL DNase I (WAKO) at 34°C for 10 min. Next, tubules were washed again with ice-cold DMEM and digested with TrypLE™ Express Enzyme (Thermo Fisher) containing 0.5 mg/mL collagenase type IV, 20 µg/mL DNase I, and 1 mM CaCl2 at 34°C for 15 min. Digestion was stopped by adding 10% fetal bovine serum (FBS; Thermo Fisher), and the single-cell suspension was filtered through a 70 µm mesh to remove somatic cells. To identify the developmental stage of each germ cell as previously described (58), cells were stained in DMEM containing 10% FBS and Hoechst 33342 (6 µg per 1 × 10^6 cells) at 34°C for 10 min.

Undifferentiated and differentiated spermatogonia were isolated using a modified protocol that combined features of two previously described methods (58, 59). Cells were stained in FACS buffer (PBS supplemented with 0.5% BSA and 2 mM EDTA) containing FITC anti-mouse CD9 antibody (BioLegend), PE anti-mouse/human CD324 (E-Cadherin) antibody (BioLegend), and Alexa Fluor® 647 anti-mouse CD117 (c-Kit) antibody (BioLegend) at 4°C for 60 min. Finally, the stained Testicular germ cells suspension was washed with FBS/DMEM and filtered again. Cells were sorted with FACS Aria III (BD) based on Hoechst Blue, Hoechst Red signals, and cell size as previously described (58).

### Isolation of spermatozoa from the caudal epididymis

Cauda epididymides were dissected from 9-week-old mice, and the surrounding adipose tissue was removed under a stereomicroscope. Each caudal epididymis was placed in a droplet of DMEM supplemented with 50 mM HEPES-KOH (pH 7.4) in a non-coated dish and finely minced with scissors to release spermatozoa. The tissue suspension was incubated at 37 °C for 30 min to allow spermatozoa to disperse into the medium. The sperm-containing medium was then collected into low-retention tubes. To assess fertility in FS and FD mice, sperm numbers were counted using a hemocytometer.

### Immunostaining of testicular germ cells

Some testes from 9–10-week-old mice were fixed in PBS containing 4% paraformaldehyde and 0.1% Triton X-100 at 4 °C for 48 h, embedded in paraffin, sectioned at 7 µm thickness, and mounted on adhesive glass slides. After deparaffinization in xylene and rehydration through a graded ethanol series, antigen retrieval was performed by heating the sections in Target Retrieval Solution (DAKO, S1699) at 120 °C for 10 min (heat-induced epitope retrieval, HIER). After gradual cooling and washing with PBS, sections were incubated with Blocking One Histo (Nacalai Tesque, cat#06349-64) at 4 °C for 1 h to prevent nonspecific antibody binding.

Paraffin sections and germ cell suspensions were then washed with PBST and incubated with primary antibodies at 4 °C overnight. Signals were detected using secondary antibodies conjugated to Alexa Fluor 488, Alexa Fluor 555, or Alexa Fluor 647 (Thermo Fisher Scientific). Nuclei were counterstained with DAPI. Fluorescence images were acquired using a confocal laser scanning microscope (LSM900; Carl Zeiss) and processed using ImageJ (National Institutes of Health).

Antibodies used in this study included: anti-H1t (kindly provided by Dr. Mary Ann Handel), anti-γH2AX (Millipore), anti-H3K4me3 (#9751; Cell Signaling Technology), anti-H3 (MABI), anti-H3K27me3 (Diagenode; C15410195), and anti-H3K9me2 (Abcam; ab1220).

### Metabolite extraction and metabolomic analysis

Metabolites were extracted from sorted germ cells as previously described (60). Briefly, cell pellets were subjected to cold organic solvent extraction to rapidly quench cellular metabolism, followed by centrifugation to remove insoluble material. The resulting extracts were analyzed using ion chromatography– mass spectrometry (IC–MS) and liquid chromatography–mass spectrometry (LC–MS). IC–MS analysis for anionic metabolites was performed using a Dionex ICS-5000+ ion chromatography system coupled to a Q Exactive Orbitrap mass spectrometer (Thermo Fisher Scientific). Metabolites were separated on an IonPac AS11-HC column using a potassium hydroxide gradient and detected in negative electrospray ionization mode as previously described (61). For cationic metabolites, LC–MS analysis was performed using an Agilent 1290 Infinity LC system coupled to a Q Exactive Orbitrap mass spectrometer. Metabolites were separated on a HILIC-Z column and detected in positive electrospray ionization mode.

Metabolite abundances were normalized to germ cell volume to account for differences in cell size across spermatogenic stages. Cell volumes were estimated based on stereological measurements of germ cell nuclear diameter and cytoplasmic proportion as described by Auharek and de França (62). Metabolites detected in at least two of the three biological replicates were retained for downstream analysis. A total of 119 metabolites met this criterion and were included in subsequent analyses. Principal component analysis (PCA) was performed to assess global metabolic differences among germ cell populations. For visualization of metabolite abundance patterns, hierarchical clustering was performed using the Morpheus web tool (Broad Institute). Clustering was conducted using one minus Pearson correlation as the distance metric with average linkage. The clustered data were visualized as a heatmap after row-wise Z-score transformation. Differential metabolite analysis between Diff and PD cells was performed using a two-sided Student’s t-test. Log_2_ fold change values were calculated as the ratio of metabolite abundance in PD relative to Diff. Volcano plots were generated to visualize differential metabolites, with the x-axis representing log_2_ fold change and the y-axis representing −log_10_ P-value. Selected metabolites belonging to representative metabolic pathways were annotated on the plot. Metabolite set enrichment analysis (MSEA) and metabolic pathway analysis were performed using MetaboAnalyst 6.0 (https://www.metaboanalyst.ca) (63).

### Protein extraction and proteomic analysis

Sorted cells were washed with cold PBS and processed using a modified surfactant-aided one-pot (SOP) protocol (64). Proteins were extracted in 0.1% n-dodecyl-β-D-maltoside (DDM) in 50 mM Tris-HCl buffer (pH 8.0) by repeated freeze–thaw cycles followed by sonication. Proteins were sequentially reduced with dithiothreitol (DTT; 37 °C, 30 min), alkylated with iodoacetamide (IAA; 37 °C, 30 min, in the dark), pre-digested with Lys-C (37 °C, 3 h), and digested with trypsin (37 °C, 16 h).

Peptide mixtures were analyzed by nanoLC–MS/MS using a nanoElute2 system coupled to a timsTOF Pro 2 mass spectrometer (Bruker) operated in diaPASEF mode (65). Raw mass spectrometry data were processed using DIA-NN (v1.8.1) for protein identification and label-free quantification against the UniProt/SwissProt mouse proteome database combined with the cRAP contaminant database (66). Three biological replicates were analyzed for each condition.

### Omni-ATAC of sorted germ cells

Omni-ATAC was performed as described previously (29, 67, 68), originally based on Buenrostro et al (37). 50,000 sorted testicular germ cells were washed in ice-cold PBS and permeabilized in 50 µL of Resuspension Buffer (10 mM Tris-HCl, pH 7.5, 10 mM NaCl, 3 mM MgCl2) containing 0.1% NP-40, 0.1% Tween-20, and 0.01% digitonin at room temperature for 30 sec. Cells were washed in Resuspension Buffer containing 0.1% Tween-20. Tagmentation was performed in 50 µL Tagmentation Buffer (10× Tn5 Transposase prepared in-house, 20 mM Tris-HCl pH 7.4, 10 mM MgCl2, 20% dimethylformamide, 0.1% Tween-20, 0.01% digitonin, 20% glycerol, 0.1% Triton X-100, and 0.33× PBS) at 37°C for 33 min with gentle resuspension. DNA was isolated as described in the library preparation section.

### CUT&Tag of sorted germ cells

CUT&Tag with optional fixation and magnetic-beads-based capture was performed as described previously (46). Briefly, 200,000 diploid cells and 1,000,000 haploid cells were washed in ice-cold PBS and permeabilized in 500 µL NE1 Buffer (20% glycerol, 0.1% Triton X-100, 0.5 mM spermidine, 10 mM KCl, 20 mM HEPES-KOH; pH 7.9) at 4°C for 30 sec. Cells were washed in ice-cold PBS and fixed in PBS containing 0.1% paraformaldehyde at room temperature for 2 min. Optionally, cells were captured on 10 µL ConA-Dynabeads, composed of Concanavalin A biotin conjugate, Type IV (Sigma), and MyOne T1 Dynabeads (Thermo Fisher), prepared in Preparation Buffer (0.01% Tween-20, 1 mM CaCl2, 1 mM MnCl2) and activated in Binding Buffer (10 mM KCl, 1 mM CaCl2, 1 mM MnCl2). Primary immunoreaction was performed with anti-H3K4me3 (#9751; Cell Signaling Technology) and anti-H3K27me3 (Diagenode; C15410195) antibody diluted in Dig-Wash Buffer (0.05% digitonin, 0.5 mM spermidine, 150 mM NaCl, 150 mM HEPES-NaOH pH 7.5, 1 mM PMSF, protease inhibitor) containing 0.1% BSA and 2 mM EDTA, followed by secondary antibody diluted in Dig-Wash Buffer. Tagmentation was initiated by binding 0.5 µL pA-Tn5 transposase prepared in-house in Dig-300 Buffer (Dig-Wash Buffer with 300 mM NaCl), started by resuspension in Dig-300 Buffer containing 10 mM MgCl2, and stopped by addition of 16.7 mM EDTA, 0.1% SDS, and 0.167 mg/mL Proteinase K after incubation at 37°C for 60 min. DNA was isolated as described below.

### Library preparation for CUT&Tag and ATAC-seq

After tagmentation in Omni-ATAC and the CUT&Tag process, genomic DNA was isolated using FastGene™ Gel/PCR Extraction Kit (Nippon Genetics), eluted in 20 µL GP3 preheated to 70°C. DNA pool was amplified by 12–14 cycles of PCR using primers based on Illumina UDP Indexes (Illumina). DNA fragments of 300–1,000 bp were size-selected using Solid Phase Reversible Immobilization (SPRI) beads prepared in-house (69) prepared in-house. Libraries were sequenced using paired-end 150-bp reads on the NovaSeq X Plus platform. Each group consisted of at least 2 biological replicates. Testes from 4–5 mice (8–10 testes per replicate) were pooled for each replicate.

### Data processing of ATAC-seq and CUT&Tag

Data analysis of ATAC-seq and CUT&Tag followed the ATAC-seq Data Standards and Processing Pipeline of the ENCODE project.

Raw paired-end reads were trimmed to remove adapters and low-quality bases using Trimmomatic v0.39 (parameters: LEADING:20, TRAILING:20, SLIDINGWINDOW:4:20, MINLEN:36) (70). Trimmed reads were aligned to the mm10 reference genome (UCSC, 2011) using Bowtie2 v2.4.1 with the parameters --end-to-end --very-sensitive --no-mixed --no-discordant -X 2000 for paired-end libraries (71). SAM files were converted to BAM, sorted, and low-quality reads (MAPQ < 30) as well as mitochondrial reads (chrM) were removed using Samtools v1.6 (72). BAM files were further filtered for properly paired reads (for paired-end libraries) and indexed. PCR duplicates were removed using Picard v2.27.5 (MarkDuplicates). For ATAC-seq, additional processing included fixmate adjustment, conversion to BEDPE as previously described (73), using Samtools and Bedtools (74). Later, Tn5 insertion site correction using bedpeTn5shift.sh, and generation of minimal BEDPE files using bedpeMinimalConvert.sh (73).

CUT&Tag libraries (H3K4me3 and H3K27me3) were processed similarly using Trimmomatic and Bowtie2, applying narrow or broad peak parameters depending on the histone mark. Filtered BAM files were indexed and duplicates removed as described above.

For quality control and visualization, BAM files were converted to BigWig coverage tracks using deepTools v3.5.2 (bamCoverage) (75) and normalized to counts per million mapped reads (CPM). Replicates were merged by first averaging coverage in bedGraph format using bigWigMerge, dividing by the number of replicates, sorting, and converting back to BigWig.

### Correlation and similarity analysis of ATAC-seq and CUT&Tag data

To assess the reproducibility and global similarity of ATAC-seq and CUT&Tag data, Pearson correlation coefficients were calculated across genomic bins for each sample. BigWig files were summarized using multiBigwigSummary from deepTools (75) with 1-kb bins across the whole genome. The resulting matrices were imported into R via the reticulate package, and CPM values were used to calculate Pearson correlations, which were visualized as heatmaps with numeric overlays using pheatmap.

For dimensionality reduction and visualization of sample relationships, the top 2,000 most variable bins were selected based on variance. Principal component analysis (PCA) was performed on log2-transformed CPM values using the prcomp function in R. All analyses were conducted in R (v4.x) using the packages ggplot2, ggrepel, pheatmap, RColorBrewer, and reticulate for integration with Python-based summary matrices.

### Peak calling for ATAC-seq and CUT&Tag

For ATAC-seq and CUT&Tag H3K4me3, narrow peaks were called using MACS2 v2.2.7.1 (76). Individual replicates were processed separately, and appropriate thresholds (ATAC: -p 1e-3; H3K4me3: -q 0.01, with --nomodel and --extsize 147) were used. Replicate peaks were merged and filtered using Bedtools to remove blacklist regions (77), and IDR (Irreproducible Discovery Rate) analysis (78) was performed to generate high-confidence, replicate-consensus peaks. Finally, union peak sets across conditions were created for downstream differential analysis.

For H3K27me3, broad peaks were called with MACS2 using --broad --broad-cutoff 0.1 -q 0.05, enabling optimal peak calling for diffuse enrichment (79). Individual replicate peaks were merged and filtered similarly to narrow peaks.

### Differential analysis for ATAC-seq and CUT&Tag

Differential analysis of chromatin accessibility and H3K4me3 was performed following a previously reported approach (73), in which union peaks were used as consensus regions and library-size normalization was applied using a loess-based method (as described in Analysis IV). Low-abundance peaks were filtered using the *csaw* package (80) based on log-CPM thresholds (logCPM ≤ −3 for ATAC-seq and logCPM ≤ 1 for CUT&Tag). Differential testing was carried out using quasi-likelihood F-tests in the *edgeR* package (81). Peaks were classified as significantly up-regulated or down-regulated and met the significance criteria (FDR < 0.05 for ATAC-seq; p < 0.05 for CUT&Tag).

For H3K27me3 CUT&Tag data, counts were transformed using the variance-stabilizing transformation (VST) implemented in *DESeq2* (82), and surrogate variable analysis (SVA) (83) was applied to adjust for potential hidden batch effects. In *DESeq2*, statistical significance was assessed using the Wald test.

### Annotation and functional enrichment analysisis for ATAC-seq and CUT&Tag

Differentially accessible or enriched regions were annotated using *HOMER* (v4.10) annotatePeaks.pl (40) with the mouse genome (mm10). Downstream analyses were performed in R (v4.x) using the *GenomicRanges, dplyr, ggplot2*, and *reshape2* packages. Genomic annotation was simplified into five categories: promoter-TSS, exon, intron, intergenic, and distances to the nearest transcription start site (TSS) were binned into ranges (0–0.2 kb, 0.2–2 kb, 2–20 kb, >20 kb).

For functional enrichment analysis, genomic regions associated with differentially accessible regions (DARs) were linked to nearby genes using the basal-plus-extension model (5 kb upstream, 1 kb downstream, with a maximum extension of 1 Mb) implemented in the *GREAT* online tool (39). Gene Ontology (GO) biological process terms enriched to nearby genes, and significance was assessed using both binomial and hypergeometric tests.

To examine chromatin compartmentalization, H3K4me3 peaks were overlapped with A/B compartment data derived from Hi-C (32, 33) at 50-kb resolution. Peaks overlapping regions with positive or negative PCA1 scores were assigned to compartment A or B.

### Motif analysis and integration with gene expression

Transcription factor (TF) motif enrichment analysis was performed on differential regions using HOMER (v4.10) findMotifsGenome.pl with the mouse mm10 genome. The resulting motif enrichment data were processed and visualized in R (v4.x) using the dplyr, readxl, ggplot2, and pheatmap packages. Top enriched motifs were extracted and cross-referenced with candidate TF names and gene expression data (TPM values from RNA-seq), which were examined across spermatogenic cell types. Only TFs that were both matched and highly expressed were included in the visualization. Subfamilies of proteins with the same target sequence were merged for clarity.

### Enrichment profiling of chromatin accessibility and histone modification

To assess enrichment of ATAC-seq and CUT&Tag around genomic regions of interest, signal matrices were computed using deepTools computeMatrix (75) in reference-point mode, centering on the midpoint of each region with ±2 kb flanking windows. The resulting matrices were visualized as heatmaps and average profiles using plotHeatmap and plotProfile, respectively.

For quantitative comparisons between experimental groups, signal intensities from bigWig files were extracted over genomic regions of interest, including DARs or differentially enriched regions, or previously defined bivalent domains (28). Signals were log10-transformed prior to visualization. Violin and box plots were generated to compare conditions, and statistical significance was evaluated using Wilcoxon rank-sum tests.

Dynamic trajectories of differentially accessible regions (DARs) were assessed by averaging bigWig signals over up- and down-regulated DARs. Log2 fold changes were then calculated, and per-region trajectories were plotted to visualize chromatin accessibility changes associated with folate deficiency.

All processing and visualization were performed in R (v4.x) using the GenomicRanges, rtracklayer, dplyr, tidyr, and ggplot2 packages.2.

## Acknowledgments

We thank Jasmine M. Esparza and Yasuhisa Munakata, and members of the S.M. laboratory for valuable discussions and helpful comments. We thank Yuka Kitamura for sharing the germ cell Hi-C dataset.A part of this work was supported by “Advanced Research Infrastructure for Materials and Nanotechnology in Japan (ARIM)” of the Ministry of Education, Culture, Sports, Science and Technology (MEXT). This work was supported by the following funding sources: Grant-in-Aid for Transformative Research Areas (A) (23H04950 to Y.H.); Grant-in-Aid for Challenging Research (Exploratory) (23K18084 to S.M.); Grant-in-Aid for Scientific Research (B) (21H02383 to S.M.); NIH grant R35GM141085 (to S.H.N.); and research grants from the TERUMO LIFE SCIENCE FOUNDATION, the Astellas Foundation for Research on Metabolic Disorders, the Takeda Science Foundation, The Naito Foundation, the Daiichi Sankyo Foundation of Life Science, the Chugai Foundation for Innovative Drug Discovery Science, and the Mochida Memorial Foundation for Medical and Pharmaceutical Research (to S.M.).

## Author contribution

I.A. and S.M. designed the study. I.A., N.F., M.M., A.H., Y.Y., T.N., and T.S. performed the experiments. I.A., N.F., M.M., A.H., and K.O. performed the computational analyses. I.A., T.S., S.H.N., Y.H., and S.M. interpreted the results. I.A., S.H.N., and S.M. wrote the manuscript with critical feedback from all authors. S.M. supervised the project.

## Competing interests

The authors declare no competing interests.

## Notes

### Competing Interest Statement

The authors have declared no competing interest.

## Reference

1. G. Cavalli, E. Heard, Advances in epigenetics link genetics to the environment and disease. Nature 571, 489–499 (2019).

2. O. J. Rando, Daddy Issues: Paternal Effects on Phenotype. Cell 151, 702–708 (2012).

3. Lismer, S. Kimmins, Emerging evidence that the mammalian sperm epigenome serves as a template for embryo development. Nature Communications 14, 2142 (2023).

4. W. Sun et al., Cold-induced epigenetic programming of the sperm enhances brown adipose tissue activity in the offspring. Nature Medicine 24, 1372–1383 (2018).

5. B. R. Carone et al., Paternally Induced Transgenerational Environmental Reprogramming of Metabolic Gene Expression in Mammals. Cell 143, 1084–1096 (2010).

6. K. Siklenka et al., Disruption of histone methylation in developing sperm impairs offspring health transgenerationally. Science 350, aab2006 (2015).

7. Lismer, K. Siklenka, C. Lafleur, V. Dumeaux, S. Kimmins, Sperm histone H3 lysine 4 trimethylation is altered in a genetic mouse model of transgenerational epigenetic inheritance. Nucleic Acids Research 48, 11380–11393 (2020).

8. D. J. Barker, C. Osmond, Infant mortality, childhood nutrition, and ischaemic heart disease in England and Wales. Lancet 1, 1077–1081 (1986).

9. P. D. Gluckman, M. A. Hanson, Living with the Past: Evolution, Development, and Patterns of Disease. Science 305, 1733–1736 (2004).

10. Soubry, Epigenetics as a Driver of Developmental Origins of Health and Disease: Did We Forget the Fathers? BioEssays 40, 1700113 (2018).

11. Y. Okada, K. Yamaguchi, Epigenetic modifications and reprogramming in paternal pronucleus: sperm, preimplantation embryo, and beyond. Cell Mol Life Sci 74, 1957–1967 (2017).

12. Q. Chen et al., Sperm tsRNAs contribute to intergenerational inheritance of an acquired metabolic disorder. Science 351, 397–400 (2016).

13. K. Xie et al., Epigenetic alterations in longevity regulators, reduced life span, and exacerbated aging-related pathology in old father offspring mice. Proceedings of the National Academy of Sciences 115, E2348–E2357 (2018).

14. M. E. Pepin et al., Antiretroviral therapy potentiates high-fat diet induced obesity and glucose intolerance. Molecular Metabolism 12, 48–61 (2018).

15. U. Sharma et al., Biogenesis and function of tRNA fragments during sperm maturation and fertilization in mammals. Science 351, 391–396 (2016).

16. K. Yoshida et al., ATF7-Dependent Epigenetic Changes Are Required for the Intergenerational Effect of a Paternal Low-Protein Diet. Molecular Cell 78, 445-458.e446 (2020).

17. M. Serefidou, A. V. Venkatasubramani, A. Imhof, The Impact of One Carbon Metabolism on Histone Methylation. Front Genet 10, 764 (2019).

18. R. Lambrot et al., Low paternal dietary folate alters the mouse sperm epigenome and is associated with negative pregnancy outcomes. Nat Commun 4, 2889 (2013).

19. L. Ly et al., Intergenerational impact of paternal lifetime exposures to both folic acid deficiency and supplementation on reproductive outcomes and imprinted gene methylation. Molecular Human Reproduction 23, 461–477 (2017).

20. Lismer et al., Histone H3 lysine 4 trimethylation in sperm is transmitted to the embryo and associated with diet-induced phenotypes in the offspring. Developmental Cell 56, 671-686.e676 (2021).

21. L. J. Luense et al., Comprehensive analysis of histone post-translational modifications in mouse and human male germ cells. Epigenetics & Chromatin 9, 24 (2016).

22. S. Erkek et al., Molecular determinants of nucleosome retention at CpG-rich sequences in mouse spermatozoa. Nature Structural & Molecular Biology 20, 868–875 (2013).

23. M. X. Qian et al., Acetylation-mediated proteasomal degradation of core histones during DNA repair and spermatogenesis. Cell 153, 1012–1024 (2013).

24. L. J. Luense et al., Gcn5-Mediated Histone Acetylation Governs Nucleosome Dynamics in Spermiogenesis. Developmental Cell 51, 745-758.e746 (2019).

25. M. D. Griswold, Spermatogenesis: The Commitment to Meiosis. Physiological Reviews 96, 1–17 (2015).

26. E. Baltus et al., In germ cells of mouse embryonic ovaries, the decision to enter meiosis precedes premeiotic DNA replication. Nature Genetics 38, 1430–1434 (2006).

27. K. Hasegawa et al., SCML2 establishes the male germline epigenome through regulation of histone H2A ubiquitination. Dev Cell 32, 574–588 (2015).

28. S. Maezawa et al., Polycomb protein SCML2 facilitates H3K27me3 to establish bivalent domains in the male germline. Proc Natl Acad Sci U S A 115, 4957–4962 (2018).

29. S. Maezawa, M. Yukawa, K. G. Alavattam, A. Barski, S. H. Namekawa, Dynamic reorganization of open chromatin underlies diverse transcriptomes during spermatogenesis. Nucleic Acids Res 46, 593–608 (2018).

30. Y. Hayashi, Y. Matsui, Metabolic Control of Germline Formation and Differentiation in Mammals. Sexual Development 16, 388–403 (2023).

31. Y. Hayashi et al., Control of epigenomic landscape and development of fetal male germ cells through L-serine metabolism. iScience 27, 110702 (2024).

32. Z. Lin et al., SETD1B-mediated broad H3K4me3 controls proper temporal patterns of gene expression critical for spermatid development. Cell Research 35, 345–361 (2025).

33. P. G. Reeves, F. H. Nielsen, G. C. Fahey, AIN-93 Purified Diets for Laboratory Rodents: Final Report of the American Institute of Nutrition Ad Hoc Writing Committee on the Reformulation of the AIN-76A Rodent Diet. The Journal of Nutrition 123, 1939–1951 (1993).

34. D. G. de Rooij, L. D. Russell, All you wanted to know about spermatogonia but were afraid to ask. J Androl 21, 776–798 (2000).

35. L. D. Russell, R. A. Ettlin, A. P. S. Hikim, E. D. Clegg, Histological and Histopathological Evaluation of the Testis. International Journal of Andrology 16, 83–83 (1993).

36. E. F. Oakberg, Duration of spermatogenesis in the mouse and timing of stages of the cycle of the seminiferous epithelium. American Journal of Anatomy 99, 507–516 (1956).

37. J. D. Buenrostro, P. G. Giresi, L. C. Zaba, H. Y. Chang, W. J. Greenleaf, Transposition of native chromatin for fast and sensitive epigenomic profiling of open chromatin, DNA-binding proteins and nucleosome position. Nat Methods 10, 1213–1218 (2013).

38. S. L. Klemm, Z. Shipony, W. J. Greenleaf, Chromatin accessibility and the regulatory epigenome. Nature Reviews Genetics 20, 207–220 (2019).

39. C. Y. McLean et al., GREAT improves functional interpretation of cis-regulatory regions. Nat Biotechnol 28, 495–501 (2010).

40. S. Heinz et al., Simple combinations of lineage-determining transcription factors prime cis-regulatory elements required for macrophage and B cell identities. Mol Cell 38, 576–589 (2010).

41. C. Yi, Y. Kitamura, S. Maezawa, S. H. Namekawa, B. R. Cairns, ZBTB16/PLZF regulates juvenile spermatogonial stem cell development through an extensive transcription factor poising network. Nature Structural & Molecular Biology 32, 1213–1226 (2025).

42. E. Lieberman-Aiden et al., Comprehensive Mapping of Long-Range Interactions Reveals Folding Principles of the Human Genome. Science 326, 289–293 (2009).

43. B. E. Bernstein et al., Genomic Maps and Comparative Analysis of Histone Modifications in Human and Mouse. Cell 120, 169–181 (2005).

44. T. Miller et al., COMPASS: A complex of proteins associated with a trithorax-related SET domain protein. Proceedings of the National Academy of Sciences 98, 12902–12907 (2001).

45. U. Sharma, O. J. Rando, Metabolic Inputs into the Epigenome. Cell Metab 25, 544–558 (2017).

46. H. S. Kaya-Okur et al., CUT&Tag for efficient epigenomic profiling of small samples and single cells. Nature Communications 10, 1930 (2019).

47. H. S. Sin, A. V. Kartashov, K. Hasegawa, A. Barski, S. H. Namekawa, Poised chromatin and bivalent domains facilitate the mitosis-to-meiosis transition in the male germline. BMC Biol 13, 53 (2015).

48. B. J. Lesch, S. J. Silber, J. R. McCarrey, D. C. Page, Parallel evolution of male germline epigenetic poising and somatic development in animals. Nat Genet 48, 888–894 (2016).

49. T. Haaf, D. C. Ward, Higher Order Nuclear Structure in Mammalian Sperm Revealed by in Situ Hybridization and Extended Chromatin Fibers. Experimental Cell Research 219, 604–611 (1995).

50. S. H. Namekawa et al., Postmeiotic Sex Chromatin in the Male Germline of Mice. Current Biology 16, 660–667 (2006).

51. S. Maezawa et al., SCML2 promotes heterochromatin organization in late spermatogenesis. J Cell Sci 131 (2018).

52. R. L. Bailey, K. P. West Jr, R. E. Black, The Epidemiology of Global Micronutrient Deficiencies. Annals of Nutrition and Metabolism 66, 22–33 (2015).

53. G. Kaati, L. O. Bygren, S. Edvinsson, Cardiovascular and diabetes mortality determined by nutrition during parents’ and grandparents’ slow growth period. European Journal of Human Genetics 10, 682–688 (2002).

54. Y. Kitamura, S. H. Namekawa, Epigenetic priming in the male germline. Current Opinion in Genetics & Development 86, 102190 (2024).

55. K. Skvortsova, N. Iovino, O. Bogdanović, Functions and mechanisms of epigenetic inheritance in animals. Nat Rev Mol Cell Biol 19, 774–790 (2018).

56. B. del Blanco et al., Kdm1a safeguards the topological boundaries of PRC2-repressed genes and prevents aging-related euchromatinization in neurons. Nature Communications 15, 1781 (2024).

57. Sakashita et al., Polycomb protein SCML2 mediates paternal epigenetic inheritance through sperm chromatin. Nucleic Acids Research 51, 6668–6683 (2023).

58. V. Gaysinskaya, I. Y. Soh, G. W. van der Heijden, A. Bortvin, Optimized flow cytometry isolation of murine spermatocytes. Cytometry A 85, 556–565 (2014).

59. T. Nakagawa et al., A multistate stem cell dynamics maintains homeostasis in mouse spermatogenesis. Cell Reports 37 (2021).

60. R. Hayasaka et al., Metabolomics of small extracellular vesicles derived from isocitrate dehydrogenase 1-mutant HCT116 cells collected by semi-automated size exclusion chromatography. Front Mol Biosci 9, 1049402 (2022).

61. Hirayama et al., The use of a double coaxial electrospray ionization sprayer improves the peak resolutions of anionic metabolites in capillary ion chromatography-mass spectrometry. Journal of Chromatography A 1619, 460914 (2020).

62. S. A. Auharek, L.R. de França, Postnatal testis development, Sertoli cell proliferation and number of different spermatogonial types in C57BL/6J mice made transiently hypo- and hyperthyroidic during the neonatal period. J Anat 216, 577–588 (2010).

63. Z. Pang et al., MetaboAnalyst 6.0: towards a unified platform for metabolomics data processing, analysis and interpretation. Nucleic Acids Research 52, W398–W406 (2024).

64. K. Martin et al., Facile One-Pot Nanoproteomics for Label-Free Proteome Profiling of 50–1000 Mammalian Cells. Journal of Proteome Research 20, 4452–4461 (2021).

65. F. Meier et al., diaPASEF: parallel accumulation–serial fragmentation combined with data-independent acquisition. Nature Methods 17, 1229–1236 (2020).

66. V. Demichev, C. B. Messner, S. I. Vernardis, K. S. Lilley, M. Ralser, DIA-NN: neural networks and interference correction enable deep proteome coverage in high throughput. Nature Methods 17, 41–44 (2020).

67. M. R. Corces et al., An improved ATAC-seq protocol reduces background and enables interrogation of frozen tissues. Nature Methods 14, 959–962 (2017).

68. M. Tatara, T. Ikeda, S. H. Namekawa, S. Maezawa, ATAC-Seq Analysis of Accessible Chromatin: From Experimental Steps to Data Analysis. Methods Mol Biol 2577, 65–81 (2023).

69. M. M. DeAngelis, D. G. Wang, T. L. Hawkins, Solid-phase reversible immobilization for the isolation of PCR products. Nucleic Acids Res 23, 4742–4743 (1995).

70. M. Bolger, M. Lohse, B. Usadel, Trimmomatic: a flexible trimmer for Illumina sequence data. Bioinformatics 30, 2114–2120 (2014).

71. B. Langmead, S. L. Salzberg, Fast gapped-read alignment with Bowtie 2. Nature Methods 9, 357–359 (2012).

72. H. Li et al., The Sequence Alignment/Map format and SAMtools. Bioinformatics 25, 2078–2079 (2009).

73. J. J. Reske, M. R. Wilson, R. L. Chandler, ATAC-seq normalization method can significantly affect differential accessibility analysis and interpretation. Epigenetics & Chromatin 13, 22 (2020).

74. R. Quinlan, I. M. Hall, BEDTools: a flexible suite of utilities for comparing genomic features. Bioinformatics 26, 841–842 (2010).

75. F. Ramírez, F. Dündar, S. Diehl, B. A. Grüning, T. Manke, deepTools: a flexible platform for exploring deep-sequencing data. Nucleic Acids Research 42, W187–W191 (2014).

76. Y. Zhang et al., Model-based Analysis of ChIP-Seq (MACS). Genome Biology 9, R137 (2008).

77. H. M. Amemiya, A. Kundaje, A. P. Boyle, The ENCODE Blacklist: Identification of Problematic Regions of the Genome. Scientific Reports 9, 9354 (2019).

78. Q. Li, J. B. Brown, H. Huang, P. J. Bickel, MEASURING REPRODUCIBILITY OF HIGH-THROUGHPUT EXPERIMENTS. The Annals of Applied Statistics 5, 1752–1779 (2011).

79. L. Abbasova et al., CUT&Tag recovers up to half of ENCODE ChIP-seq histone acetylation peaks. Nature Communications 16, 2993 (2025).

80. T. L. Lun, G. K. Smyth, csaw: a Bioconductor package for differential binding analysis of ChIP-seq data using sliding windows. Nucleic Acids Research 44, e45–e45 (2016).

81. M. D. Robinson, D. J. McCarthy, G. K. Smyth, edgeR: a Bioconductor package for differential expression analysis of digital gene expression data. Bioinformatics 26, 139–140 (2010).

82. M. I. Love, W. Huber, S. Anders, Moderated estimation of fold change and dispersion for RNA-seq data with DESeq2. Genome Biology 15, 550 (2014).

83. J. T. Leek, J. D. Storey, Capturing heterogeneity in gene expression studies by surrogate variable analysis. PLoS Genet 3, 1724–1735 (2007).

