## Supplementary Figure for "Folate-dependent one-carbon metabolism controls meiotic and post-meiotic epigenome remodeling in the male germline"

### Supplementary figure legends

**Fig. S1. Metabolic pathway enrichment analysis of metabolites increased in Diff cells.**

(A) Metabolite set enrichment analysis was performed using MetaboAnalyst 6.0 for metabolites significantly increased in Diff relative to PD (42 metabolites). The top 10 enriched metabolic pathways ranked by $-\log_{10} P$-value are shown. Circle size represents the enrichment ratio for each pathway.

**Fig. S2. Physiological and reproductive phenotypes under dietary folate manipulation.**

(A) Schematic overview of the mouse experimental design. Male mice were fed a folate-sufficient control diet or a folate-deficient diet from 3 to 9 weeks of age

(B) Composition of the experimental diets used in this study. The left table shows the percentage of each component by weight. The right table lists the composition of the vitamin mix included in the diets.

(C) Body weight changes during the feeding period. Mean body weight of mice (n = 4 per group) was monitored throughout the dietary intervention and is shown as a line plot. Statistical comparisons among groups at each time point were performed using two-sided Welch's t-tests.

(D) Testis weight normalized to body weight measured at 9 weeks of age (n = 8 per group). Statistical comparisons were performed using a two-sided Welch's t-test.

(E) Fertility test. Each experimental male (n = 4 per group) was housed with three females maintained on a standard diet to minimize potential maternal dietary effects for three consecutive days under natural mating conditions. The average number of offspring produced per male is shown. No significant differences in litter size were observed between groups (two-sided Welch's t-test).

(F) Epididymal sperm counts measured from cauda epididymides of adult mice (n = 7 per group). Statistical comparisons were performed using a two-sided Welch's t-test.

(G) Quantification of germ cell populations within seminiferous tubules based on histological analysis. Testes were collected at 9 weeks of age, fixed, paraffin-embedded, and subjected to immunofluorescence staining. Seminiferous tubule stages were classified into I–III, IV–VII, and VIII–IX based on staining patterns using antibodies against γH2AX and H1T, markers of germ cell differentiation.

(H) Distribution of major germ cell populations within seminiferous tubules. Cells were categorized into three groups corresponding to key stages of spermatogenesis: pre-meiotic cells (Type A spermatogonia to leptotene spermatocytes), pachytene spermatocytes, and round spermatids. The relative proportions of these cell populations within seminiferous tubule cross-sections are shown as bar graphs.

**Fig. S3. Quality assessment of genome-wide ATAC-seq signal profiles.**

(A) Principal component analysis (PCA) of ATAC-seq signals in PD and RS. CPM-normalized bigWig files were summarized across the mouse genome (mm10) using 1-kb bins with the multiBigwigSummary function from deepTools, generating the same bin-based matrix used for PCA analysis and the Pearson correlation heatmap. The matrix was log2-transformed (log2[CPM + 1]), and the top 2,000 most variable bins across samples were selected for PCA. Principal components were calculated using the prcomp function in R with scaling enabled. Each point represents an individual sample, colored according to folate condition (FS or FD), and ellipses indicate the 75% confidence interval for each group.

(B) Pearson correlation heatmap of ATAC-seq signals in Diff, PD, and RS. The resulting matrix was log2-transformed (log2[CPM+1]), and pairwise Pearson correlation coefficients were calculated between samples. Correlation values are shown in each cell of the heatmap.

**Fig. S4. Functional annotation and transcription factor analysis of differential chromatin accessibility.**

(A) Distribution of distances to TSSs for upDARs, downDARs, and non-significant regions in Diff, PD, and RS. Regions were categorized according to their distance from the nearest RefSeq-annotated TSS. Bars represent the percentage of regions within each distance category.

(B) Putative transcription factors (TFs) enriched in differentially enriched regions of DARs in Diff are shown as a balloon plot. DNA sequence logos represent TF-binding motifs identified using the HOMER package. Bubble size indicates the percentage of target sequences containing the motif, and color represents the –log10 P-value. Germ cell–expressed transcription factors among the top enriched motifs are highlighted and shown with their expression levels (log2 TPM) across spermatogenic stages in the accompanying heatmap.

(C) Gene Ontology (GO) biological process terms enriched among genes associated with DARs in PD and RS. Genomic regions were linked to genes using the basal-plus-extension model (5 kb upstream, 1 kb downstream, with extension up to 1 Mb) implemented in the GREAT online analyzer. The significance of each GO term was evaluated using both hypergeometric and binomial tests and ranked based on P-values from both tests. Bar length represents the significance (–log10 binomial P-value) of each selected term.

**Fig. S5. H3K4me3 chromatin landscapes in round spermatids.**

(A) Pearson correlation heatmap of genome-wide H3K4me3 CUT&Tag signals in RS. CPM-normalized bigWig files were summarized across the mouse genome (mm10) using 1-kb bins with the multiBigwigSummary function from deepTools. The resulting matrix was log2-transformed (log2[CPM+1]), and pairwise Pearson correlation coefficients were calculated between samples. Correlation values are shown in each cell of the heatmap.

(B) Putative transcription factors (TFs) enriched in differentially enriched regions of H3K4me3 in RS are shown as a balloon plot. DNA sequence logos represent TF-binding motifs identified using the HOMER package. Bubble size indicates the percentage of target sequences containing the motif, and color represents the –log10 P-value. Germ cell–expressed transcription factors among the top enriched motifs are highlighted and shown with their expression levels (log2 TPM) across spermatogenic stages in the accompanying heatmap.

**Fig. S6. Genome-wide analysis of H3K27me3 dynamics in meiotic cells.**

(A) Pearson correlation heatmap of genome-wide H3K27me3 CUT&Tag signals in PD. CPM-normalized bigWig files were summarized across the mouse genome (mm10) using 1-kb bins with the multiBigwigSummary function from deepTools. The resulting matrix was log2-transformed (log2[CPM+1]), and pairwise Pearson correlation coefficients were calculated between samples. Correlation values are shown in each cell of the heatmap.

(B) Pearson correlation heatmap of genome-wide H3K27me3 CUT&Tag signals in round spermatids (RS), calculated as described in (A).

(C) MA plot showing differential H3K27me3 enrichment between FS and FD cells in PD. Each dot represents a broad H3K27me3 peak region derived from the consensus peak set. The x-axis indicates the average of normalized raw counts (baseMean) calculated by DESeq2, and the y-axis shows the log2 fold change (FD vs. FS). The dashed horizontal line denotes no change (log2FC = 0). Regions with $P < 0.05$ were defined as differentially enriched, with increased H3K27me3 levels in FD (upH3K27me3; red) and decreased levels in FD (downH3K27me3; blue), while non-significant regions are shown in gray. Differential analysis was performed using DESeq2 on raw count matrices derived from broad peak regions, incorporating size-factor normalization and surrogate variable adjustment to account for latent batch effects.

(D) Distribution of genomic annotations for upH3K27me3, downH3K27me3, and not significant regions in PD. Peaks were categorized into promoter–TSS, exon, intron, intergenic, and other regions based on RefSeq gene annotations. Bars represent the percentage of regions within each category, consistent with the broad genomic distribution characteristic of H3K27me3 domains.

(E) Average tag density profiles of H3K4me3, H3K27me3 CUT&Tag, and ATAC-seq signals in RS centered on TSS of previously reported bivalent genes. Signal intensity was calculated across ±2 kb from the TSS using a reference-point approach. The y-axis represents average signal intensity.

(F) Violin plots showing the distribution of H3K4me3, H3K27me3 CUT&Tag, and ATAC-seq enrichment in RS at TSS of developmental genes. The average signal was quantified and compared between folate-sufficient (FS) and folate-deficient (FD) RS. Signal intensity is shown as $\log_{10}$-transformed normalized coverage with a pseudocount added ($\log_{10}$[signal + 0.01]) to avoid zero values. Violin plots represent distribution density, with overlaid boxplots indicating the median and interquartile range. Statistical significance between FS and FD was assessed using a two-sided Wilcoxon rank-sum test and is indicated using standard notation (*P < 0.05, **P < 0.01, ***P < 0.001, n.s., not significant).

#### Fig. S7 Genome browser views of sperm H3K4me3 changes.

Genome browser tracks displaying sperm H3K4me3 ChIP-seq signals under FS (yellow) and FD (pink) conditions, together with H3K4me3 CUT&Tag signals in RS to contextualize the dynamics of H3K4me3 during spermiogenesis and sperm maturation. Signals are shown as CPM-normalized coverage. Regions highlighted in red and blue correspond to upH3K4me3 and downH3K4me3 in sperm, respectively. RefSeq gene models are shown at the top, and the black bar indicates genomic scale.

**Figure S1**

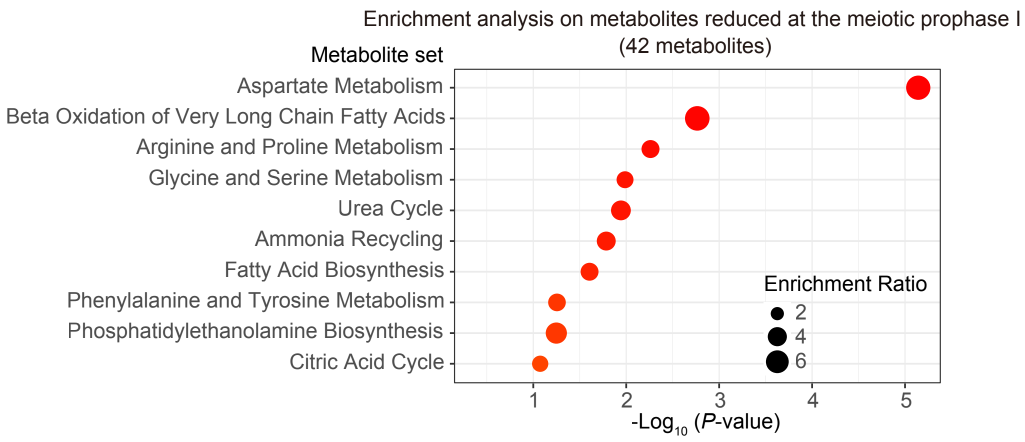
Enrichment analysis on metabolites reduced at the meiotic prophase I
(42 metabolites)
Metabolite set
Aspartate Metabolism
Beta Oxidation of Very Long Chain Fatty Acids
Arginine and Proline Metabolism
Glycine and Serine Metabolism
Urea Cycle
Ammonia Recycling
Fatty Acid Biosynthesis
Phenylalanine and Tyrosine Metabolism
Phosphatidylethanolamine Biosynthesis
Citric Acid Cycle
Enrichment Ratio
2
4
6
1
2
3
4
5
-Log10 (P-value)

**Figure S2**

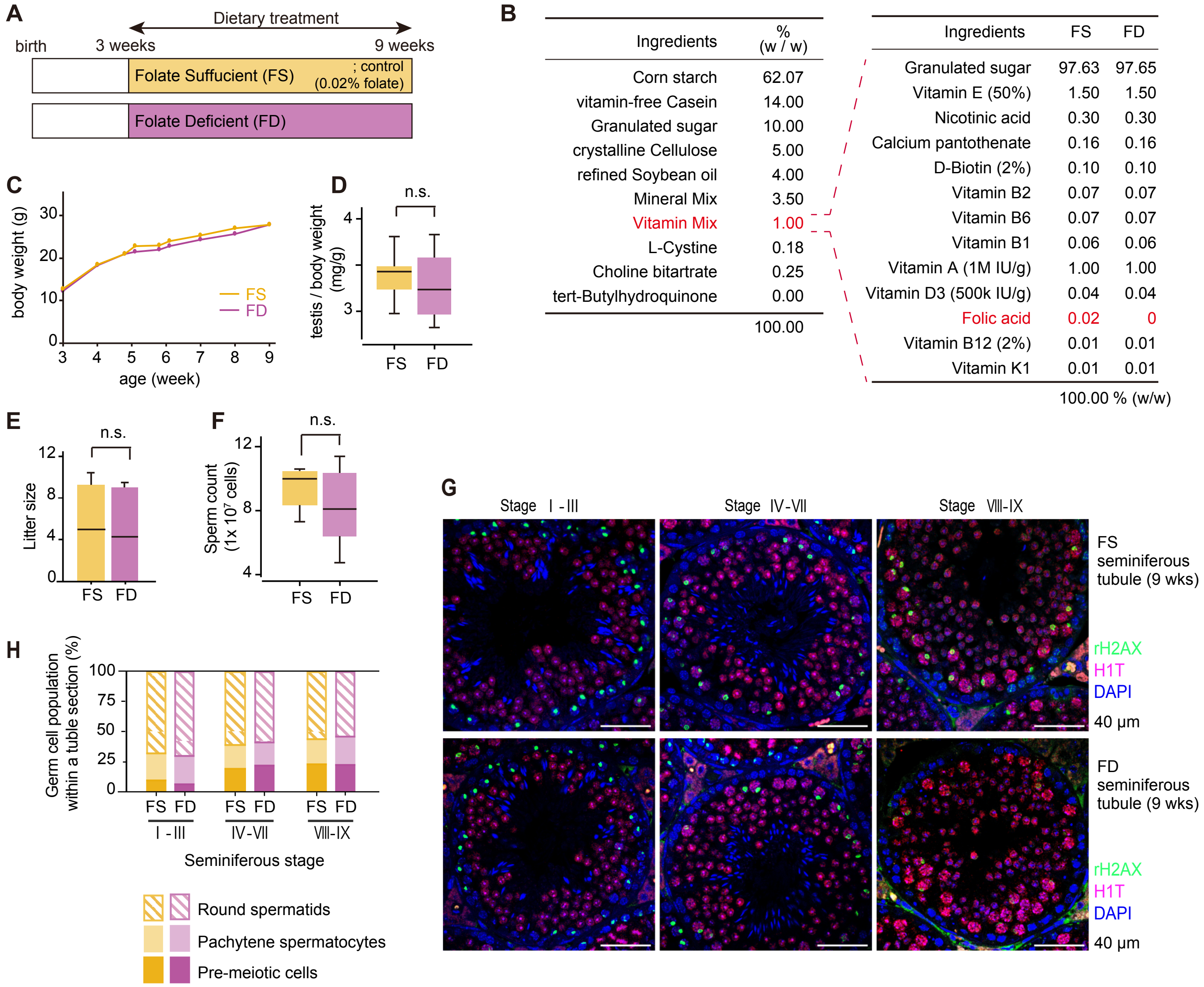


| Ingredients | % (w / w) |
|---|---|
| Corn starch | 62.07 |
| vitamin-free Casein | 14.00 |
| Granulated sugar | 10.00 |
| crystalline Cellulose | 5.00 |
| refined Soybean oil | 4.00 |
| Mineral Mix | 3.50 |
| Vitamin Mix | 1.00 |
| L-Cystine | 0.18 |
| Choline bitartrate | 0.25 |
| tert-Butylhydroquinone | 0.00 |
| | 100.00 |

| Ingredients | FS | FD |
|---|---|---|
| Granulated sugar | 97.63 | 97.65 |
| Vitamin E (50%) | 1.50 | 1.50 |
| Nicotinic acid | 0.30 | 0.30 |
| Calcium pantothenate | 0.16 | 0.16 |
| D-Biotin (2%) | 0.10 | 0.10 |
| Vitamin B2 | 0.07 | 0.07 |
| Vitamin B6 | 0.07 | 0.07 |
| Vitamin B1 | 0.06 | 0.06 |
| Vitamin A (1M IU/g) | 1.00 | 1.00 |
| Vitamin D3 (500k IU/g) | 0.04 | 0.04 |
| Folic acid | 0.02 | 0 |
| Vitamin B12 (2%) | 0.01 | 0.01 |
| Vitamin K1 | 0.01 | 0.01 |
| | 100.00 % (w/w) | |

**Figure S3**

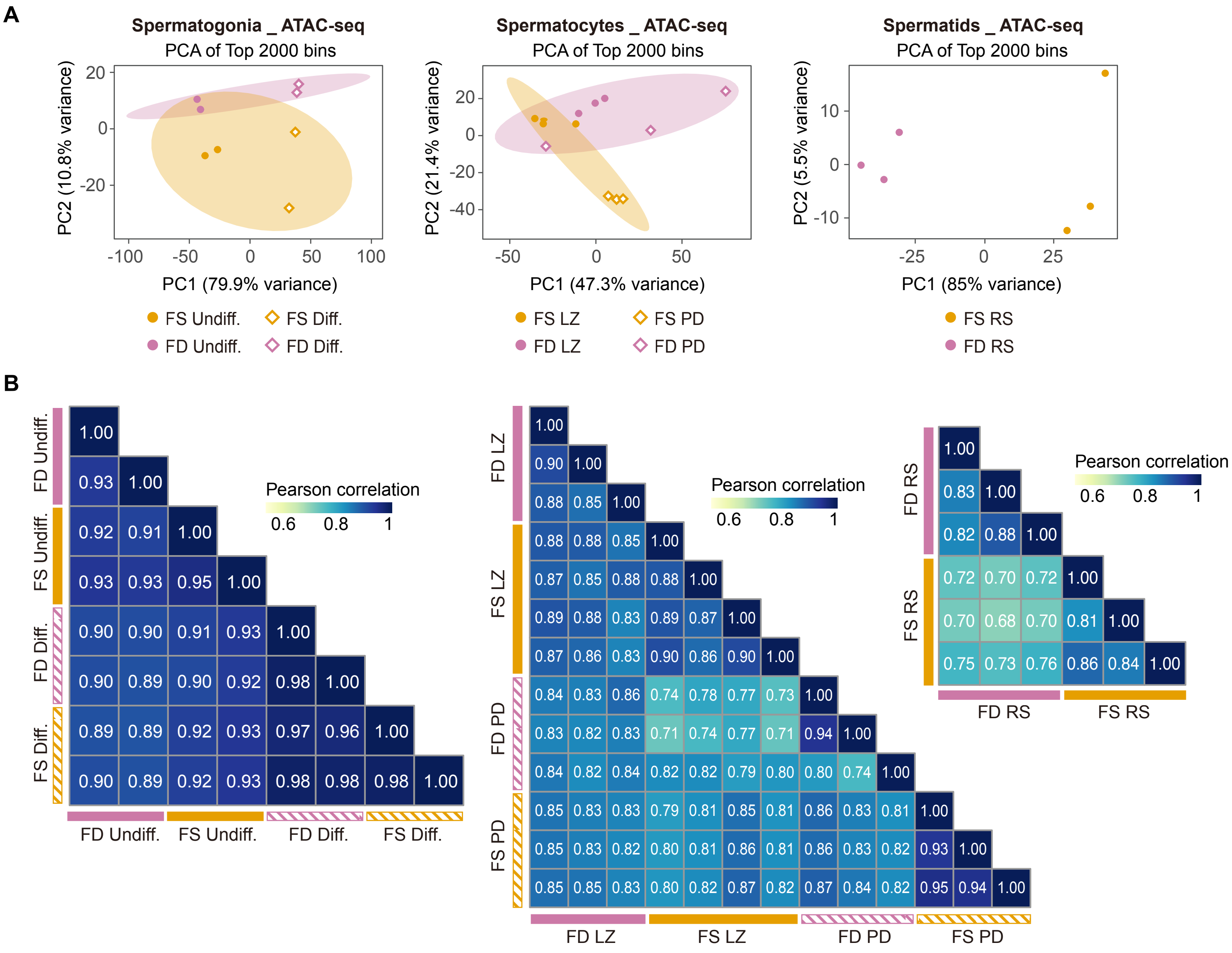
A
Spermatogonia _ ATAC-seq
PCA of Top 2000 bins
PC2 (10.8% variance)
PC1 (79.9% variance)
FS Undiff.
FS Diff.
FD Undiff.
FD Diff.
Spermatocytes _ ATAC-seq
PCA of Top 2000 bins
PC2 (21.4% variance)
PC1 (47.3% variance)
FS LZ
FS PD
FD LZ
FD PD
Spermatids _ ATAC-seq
PCA of Top 2000 bins
PC2 (5.5% variance)
PC1 (85% variance)
FS RS
FD RS
B
Pearson correlation
0.6 0.8 1
FD Undiff.
FS Undiff.
FD Diff.
FS Diff.
1.00
0.93 1.00
0.92 0.91 1.00
0.93 0.93 0.95 1.00
0.90 0.90 0.91 0.93 1.00
0.90 0.89 0.90 0.92 0.98 1.00
0.89 0.89 0.92 0.93 0.97 0.96 1.00
0.90 0.89 0.92 0.93 0.98 0.98 0.98 1.00
FD LZ
FS LZ
FD PD
FS PD
1.00
0.90 1.00
0.88 0.85 1.00
0.88 0.88 0.85 1.00
0.87 0.85 0.88 0.88 1.00
0.89 0.88 0.83 0.89 0.87 1.00
0.87 0.86 0.83 0.90 0.86 0.90 1.00
0.84 0.83 0.86 0.74 0.78 0.77 0.73 1.00
0.83 0.82 0.83 0.71 0.74 0.77 0.71 0.94 1.00
0.84 0.82 0.84 0.82 0.82 0.79 0.80 0.80 0.74 1.00
0.85 0.83 0.83 0.79 0.81 0.85 0.81 0.86 0.83 0.81 1.00
0.85 0.83 0.82 0.80 0.81 0.86 0.81 0.86 0.83 0.82 0.93 1.00
0.85 0.85 0.83 0.80 0.82 0.87 0.82 0.87 0.84 0.82 0.95 0.94 1.00
FD RS
FS RS
1.00
0.83 1.00
0.82 0.88 1.00
0.72 0.70 0.72 1.00
0.70 0.68 0.70 0.81 1.00
0.75 0.73 0.76 0.86 0.84 1.00

### Figure S4

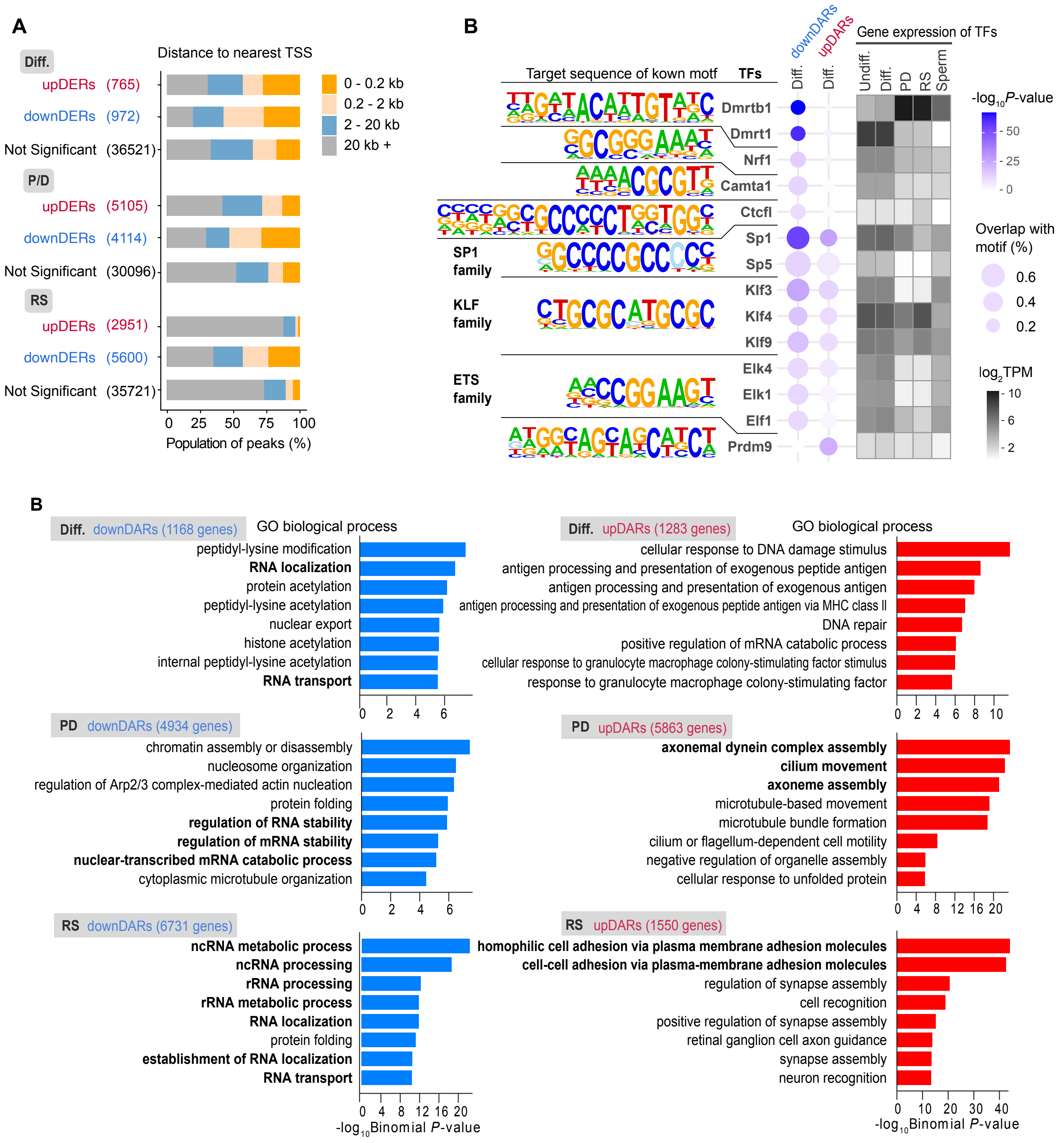
A
Distance to nearest TSS
Diff.
upDERs (765)
downDERs (972)
Not Significant (36521)
P/D
upDERs (5105)
downDERs (4114)
Not Significant (30096)
RS
upDERs (2951)
downDERs (5600)
Not Significant (35721)
0 - 0.2 kb
0.2 - 2 kb
2 - 20 kb
20 kb +
0 25 50 75 100
Population of peaks (%)
B
Target sequence of kown motf
TFs
downDARs
upDARs
Diff.
Diff.
Gene expression of TFs
Undiff.
Diff.
PD
RS
Sperm
Dmrtb1
Dmrt1
Nrf1
Camta1
Ctcfl
SP1 family
Sp1
Sp5
KLF family
Klf3
Klf4
Klf9
ETS family
Elk4
Elk1
Elf1
Prdm9
-log10P-value
50
25
0
Overlap with motif (%)
0.6
0.4
0.2
log2TPM
10
6
2
B
Diff. downDARs (1168 genes)
GO biological process
peptidyl-lysine modification
RNA localization
protein acetylation
peptidyl-lysine acetylation
nuclear export
histone acetylation
internal peptidyl-lysine acetylation
RNA transport
0 2 4 6
Diff. upDARs (1283 genes)
GO biological process
cellular response to DNA damage stimulus
antigen processing and presentation of exogenous peptide antigen
antigen processing and presentation of exogenous antigen
antigen processing and presentation of exogenous peptide antigen via MHC class II
DNA repair
positive regulation of mRNA catabolic process
cellular response to granulocyte macrophage colony-stimulating factor stimulus
response to granulocyte macrophage colony-stimulating factor
0 2 4 6 8 10
PD downDARs (4934 genes)
chromatin assembly or disassembly
nucleosome organization
regulation of Arp2/3 complex-mediated actin nucleation
protein folding
regulation of RNA stability
regulation of mRNA stability
nuclear-transcribed mRNA catabolic process
cytoplasmic microtubule organization
0 2 4 6
PD upDARs (5863 genes)
axonemal dynein complex assembly
cilium movement
axoneme assembly
microtubule-based movement
microtubule bundle formation
cilium or flagellum-dependent cell motility
negative regulation of organelle assembly
cellular response to unfolded protein
0 4 8 12 16 20
RS downDARs (6731 genes)
ncRNA metabolic process
ncRNA processing
rRNA processing
rRNA metabolic process
RNA localization
protein folding
establishment of RNA localization
RNA transport
0 4 8 12 16 20
-log10Binomial P-value
RS upDARs (1550 genes)
homophilic cell adhesion via plasma membrane adhesion molecules
cell-cell adhesion via plasma-membrane adhesion molecules
regulation of synapse assembly
cell recognition
positive regulation of synapse assembly
retinal ganglion cell axon guidance
synapse assembly
neuron recognition
0 10 20 30 40
-log10Binomial P-value

**Figure S5**

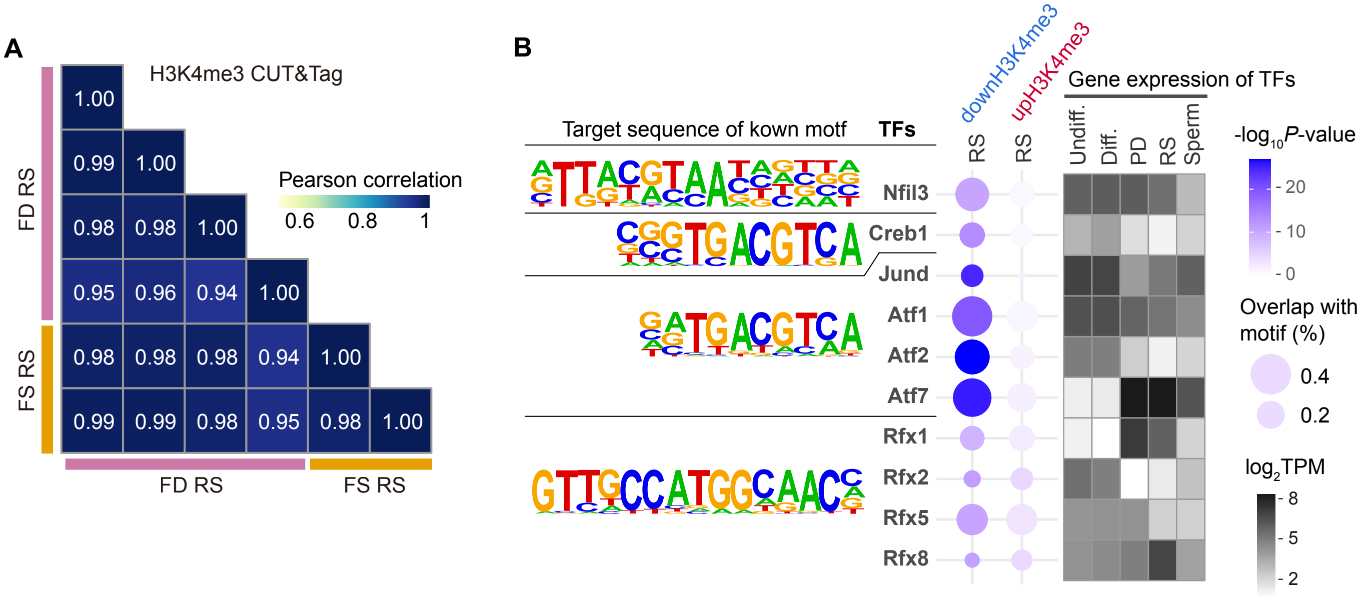
A
H3K4me3 CUT&Tag
Pearson correlation
0.6
0.8
1
FD RS
FS RS
1.00
0.99 1.00
0.98 0.98 1.00
0.95 0.96 0.94 1.00
0.98 0.98 0.98 0.94 1.00
0.99 0.99 0.98 0.95 0.98 1.00
B
Target sequence of kown motf
TFs
downH3K4me3
upH3K4me3
RS
RS
Gene expression of TFs
Undiff.
Diff.
PD
RS
Sperm
Nfil3
Creb1
Jund
Atf1
Atf2
Atf7
Rfx1
Rfx2
Rfx5
Rfx8
-log10P-value
20
10
0
Overlap with motif (%)
0.4
0.2
log2TPM
8
5
2

**Figure S6**

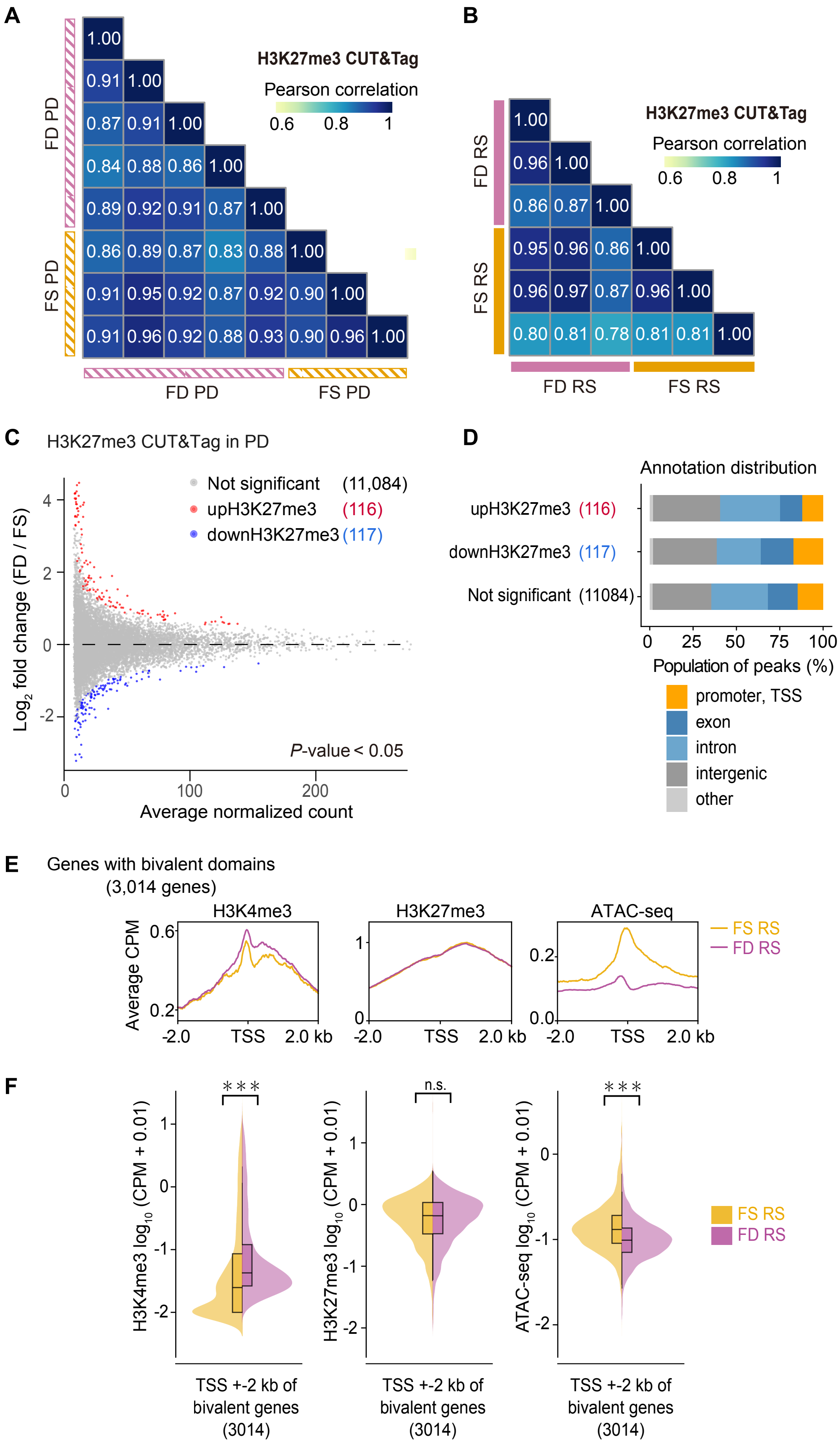
A
H3K27me3 CUT&Tag
Pearson correlation
0.6 0.8 1
FD PD
FS PD
B
H3K27me3 CUT&Tag
Pearson correlation
0.6 0.8 1
FD RS
FS RS
C
H3K27me3 CUT&Tag in PD
Not significant (11,084)
upH3K27me3 (116)
downH3K27me3 (117)
Log2 fold change (FD / FS)
P-value < 0.05
Average normalized count
D
Annotation distribution
upH3K27me3 (116)
downH3K27me3 (117)
Not significant (11084)
Population of peaks (%)
promoter, TSS
exon
intron
intergenic
other
E
Genes with bivalent domains (3,014 genes)
H3K4me3
H3K27me3
ATAC-seq
FS RS
FD RS
Average CPM
-2.0 TSS 2.0 kb
F
H3K4me3 log10 (CPM + 0.01)
H3K27me3 log10 (CPM + 0.01)
ATAC-seq log10 (CPM + 0.01)
n.s.
FS RS
FD RS
TSS +-2 kb of bivalent genes (3014)

**Figure S7**

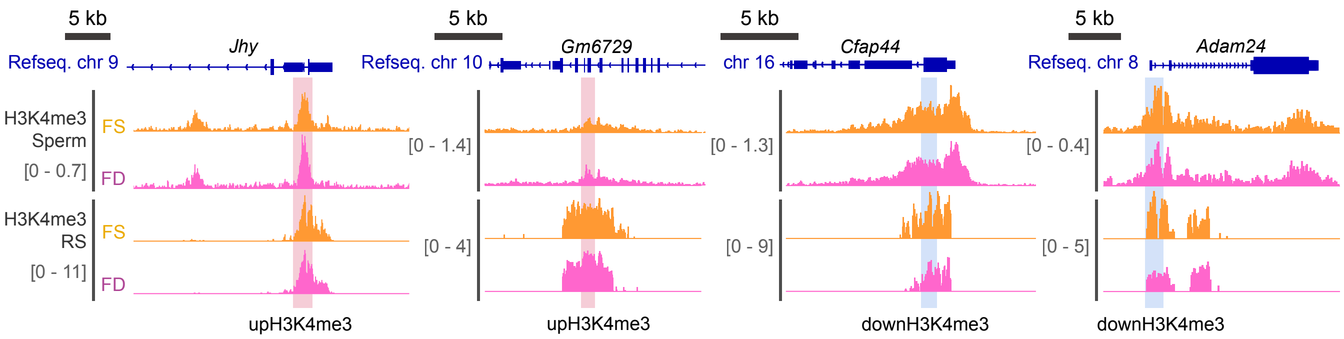
5 kb
Refseq. chr 9
Jhy
H3K4me3
Sperm
[0 - 0.7]
FS
FD
H3K4me3
RS
[0 - 11]
FS
FD
upH3K4me3
5 kb
Refseq. chr 10
Gm6729
[0 - 1.4]
[0 - 4]
upH3K4me3
5 kb
chr 16
Cfap44
[0 - 1.3]
[0 - 9]
downH3K4me3
5 kb
Refseq. chr 8
Adam24
[0 - 0.4]
[0 - 5]
downH3K4me3
